# A new view of dorsal visual cortex in macaques revealed by submillimeter fMRI

**DOI:** 10.1101/283051

**Authors:** Qi Zhu, Wim Vanduffel

## Abstract

Macaque dorsal occipital cortex is generally thought to contain one elongated third-tier area, V3d, extending along most of the rostral V2d border. In contrast, our submillimeter retinotopic fMRI maps (0.6 mm isotropic voxels, achieved by implanted phased-array receive coils) consistently show three sectors anterior to V2d. The dorsal (mirror-image) sector complies with the traditional V3d definition, and the middle (non-mirror image) sector with V3A. The ventral (mirror-image) sector bends away from V2d, as does VLP in marmosets and DLP in owl monkeys, and represents the entire contralateral hemifield like V3A. Its population receptive field size, however, suggests that this ventral sector is another fourth-tier area caudal to V4d. Hence, contrary to prevailing views, the retinotopic organization of cortex rostral to V2d differs substantially from widely accepted models. Instead, it is evolutionarily largely conserved in Old and New World monkeys.

## Introduction

The visuotopic organization of third-tier visual areas in macaque monkeys has been a longstanding source of controversy, even ∼40 years after their discovery (Angelucci and Rosa, 2015; Angelucci et al., 2015; Gattass et al., 2015; Jeffs et al., 2015; Kaas et al., 2015; Sereno et al., 2015). Different visuotopic representations have been described immediately rostral to the secondary visual area (V2), leading to a profusion of partitioning models for the third visual area V3 (Figure 1). The currently most popular model (macaque model 1, Figure 1A) proposes a V3, split into upper (UVF) and lower visual field (LVF) representations mirroring that of V2 in ventral (V3v) and dorsal V3 (V3d), respectively (see also model 3 in Figure 1E). The first evidence supporting model 1 came from connectivity (Cragg, 1969; Zeki, 1969) and single-cell receptive field mapping studies (Zeki, 1978a), and suggests a V3 extending along the entire rostral V2 border. This was modified by Colby et al. (1988), proposing that a parieto-occipital area PO [afterwards redefined as the sixth visual area (V6) by Galletti et al. (1996)], instead of V3d, borders the most dorsomedial part of V2. Later, Gattass et al. (1988) described a small gap splitting V3d and V3v, and another gap in the middle of V3d of some subjects, splitting the representation of the lower vertical meridian (LVM) at the anterior V3d border (Figure 1A, see also model 3 in Figure 1E). Similar organizational schemes have been proposed for New World monkeys (e.g. Lyon and Kaas, 2001; 2002a) and Prosimians (Fan et al., 2012; Lyon and Kaas, 2002b), although the majority of New World monkey studies favour alternative models (e.g. Figure 1B, C). V3d and V3v in macaques have also been considered distinct areas, with V3v corresponding to ventroposterior area VP, based on afferent projections from V1 present in V3d but absent in VP (Burkhalter et al., 1986; Felleman et al., 1997). In this model, V3d is highly myelinated with more direction- and fewer color-selective neurons compared to VP (Burkhalter et al., 1986; Felleman and Van Essen, 1987). This proposal, however, has been considered improbable, as both areas represent only one quadrant (Zeki, 2003). Moreover, afferent projections from V1 were also observed in V3v using more sensitive tracers (Lyon and Kaas, 2002c). Hence, the separation of V3d and V3v into independent areas is still controversial. Rostral to V3d, another third-tier area, V3A, is described, with a representation of the entire contralateral hemifield. V3A shares a LVM at its posterior border with V3d, and an upper vertical meridian (UVM) at its rostral border with the fourth visual area (V4) (Van Essen and Zeki, 1978). A similar organization was also observed by Colby et al. (1988), but redefined by Gattass et al. (1988) based on detailed electrophysiological visual field mapping, with an UVF posteriorly and a LVM bordering V4 anteriorly (Figure 1A).

**Figure 1.**
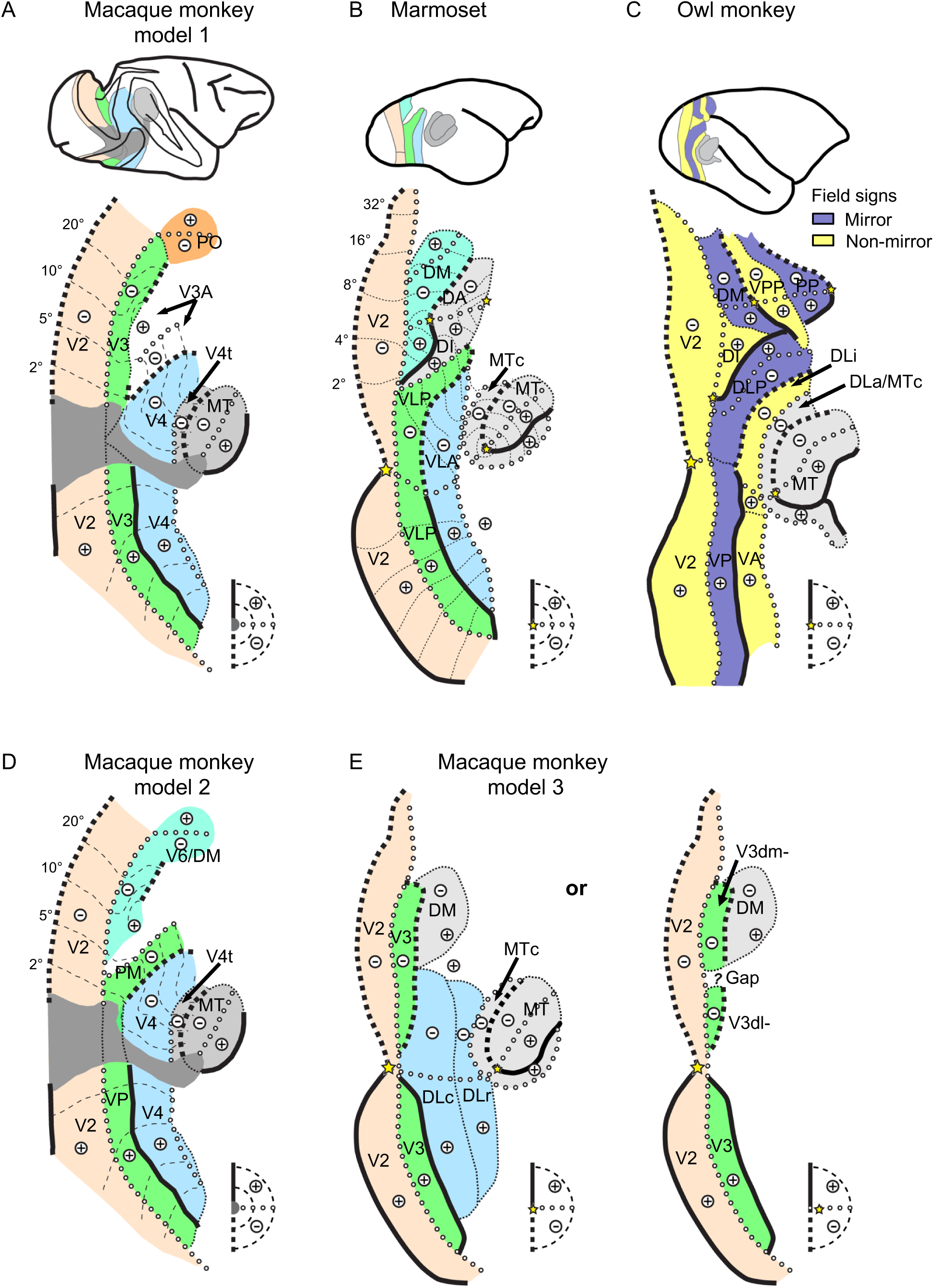
Different visuotopic models of third- and fourth-tier visual areas in Old and New World monkeys. ***A*** Most-widely accepted macaque model based on Gattass et al. (1988; 2015). ***B*** New World marmoset model based on Rosa and Tweedale (2005) and Angelucci and Rosa (2015). ***C*** New World owl monkey model based on Sereno et al. (2015). ***D*** Re-interpretation of macaque model 1 in ***A*** by Rosa and Tweedale (2005) and Angelucci and Rosa (2015) based on the marmoset model. ***E*** Alternative macaque model of Kaas and co-workers. The model on the left is based on Stepniewska et al. (2005). Thin dashed lines are drawn for the anterior borders of DLc and DLr since visual field representation is difficult to discern from the results of Stepniewska and Kaas (1996) and Stepniewska et al. (2005). The modified V3d model on the right is based on Kaas et al. (2015), in which a gap separates V3d into a dorsomedial (V3dm) and a dorsolateral (V3dl) part. V1, primary visual area; V2, second visual area; V3, third visual area; V3A, a third-tier area anterior to V3; V4, fourth visual area; V4t, transitional V4; MT, middle temporal visual area; PO, parieto-occipital area; DM, dorsomedial area; DA, dorsoanterior area; DI, dorsointermediate area; VLP, ventrolateral posterior area; VLA, ventrolateral anterior area; MTc, MT crescent; VPP, Ventral posterior parietal area; PP, Posterior parietal area; DLP, Dorsolateral posterior area; VP, Ventroposterior area; DLi, Dorsolateral intermediate area; DLa, Dorsolateral anterior area; VA, Ventroanterior area; V6, sixth visual area; PM, posteromedial area; DLc, caudal division of dorsolateral visual complex; DLr, rostral division of dorsolateral visual complex.

In another model based on New World monkeys, V3d corresponds to dorsomedial area DM (Figure 1B, C). DM in New World monkeys occupies the same topographic cortical location as macaque V3d and is also more densely myelinated compared to its neighbours. However, unlike V3d, DM represents a complete contralateral hemifield, and receives inputs from the entire extent of V1, hence it is a complete area. On the other hand, V3v in marmosets is part of a separate ventrolateral posterior region, VLP, with a LVF representation curving rostrally away from V2. VLP shares a vertical meridian (VM) anteriorly with a ventrolateral anterior region (VLA) that corresponds to macaque V4 (Figure 1B). Although not immediately apparent, this New World monkey model could be compatible with the macaque data, since the upper quadrant of V3A in macaque model 1 can be combined with the most medial part of V3d and PO/V6 to form a DM-like region as in macaque model 2 (Figure 1D). Moreover, the lower-quadrant representation assigned to V3A in macaque model 1 resembles the LVF representation of VLP (VLP-) in the New World monkey model (Figure 1D) (Angelucci and Rosa, 2015; Rosa and Tweedale, 2005).

Similar controversies exist for area V4. In the widely-accepted macaque model (model 1), V4 contains a single representation of the contralateral hemifield, with the LVF and UVF represented in dorsal (V4d) and ventral V4 (V4v), respectively (Figure 1A). Dorsally, V4d extends from the anterior bank of the lunate sulcus to the posterior bank of the superior temporal sulcus (STS). It borders V3d and V3A posteriorly and transitional V4 (V4t) anteriorly, with vertical and horizontal meridians (HM) establishing its posterior and anterior borders, respectively. This hemifield representation has also been observed in an area also named V4 in New World cebus monkeys (Piñon et al., 1998) and VLA of New World marmoset monkeys (Figure 1B) (Rosa and Tweedale, 2000b). However, other studies have suggested two parallel representations of the LVF quadrant between V3d and V4t in macaques (Baizer and Maguire, 1983; Maguire and Baizer, 1984; Youakim et al., 2001). Although in conflict with the Gattass et al. claim of a single visual field representation in V4 (Gattass et al., 1988; 2015), this is consistent with the earliest definitions of fourth-tier areas (Van Essen and Zeki, 1978; Zeki, 1971) suggesting the existence of multiple visual field representations including, for example, V4 proper and V4A in the macaque “V4 complex”. The existence of V4 and V4A was also confirmed by electrophysiology (Pigarev et al., 2002) and recent fMRI-based retinotopic mapping studies (Arcaro and Livingstone, 2017a; 2017b; Janssens et al., 2014; Kolster et al., 2014). Similarly, a V4-like dorsolateral area DL (Allman and Kaas, 1974) in New World owl monkeys has been subdivided into caudal and rostral divisions (DLc and DLr), based on connectivity patterns (Cusick and Kaas, 1988), and a separate V4t-like area corresponds to the MT crescent (MTc) (Kaas and Morel, 1993) (Figure 1E). Connectivity evidence for similar subdivisions of DL/V4 has been observed in macaque monkeys (Stepniewska and Kaas, 1996; Stepniewska et al., 2005), arguing for a more complex organization of macaque area V4 (model 3, Figure 1E). In summary, as with third-tier visual areas, the topographic organization of area V4 in macaque monkeys remains contentious.

The most likely source of these controversies relates to the incomplete evidence supporting any of these models. The sequential nature and finite sample size of tractography and microelectrode recordings may have led to misinterpretations because of uneven sampling and registration errors from 2D sections to 3D brains. Functional magnetic resonance imaging (fMRI), in contrast, has the advantage of revealing large-scale, detailed topographic information simultaneously across the entire brain. Several fMRI studies (Arcaro and Livingstone, 2017a; 2017b; Brewer et al., 2002; Fize et al., 2003; Janssens et al., 2014; Kolster et al., 2014; 2009) have acquired detailed retinotopic maps encompassing the visual cortex of macaque monkeys. The most recent results apparently confirm the widely-accepted macaque model 1. However, fine-grained information may have been missed in these studies because of their relatively low spatial resolution.

Therefore, to evaluate these models, we acquired phase-encoded retinotopic maps in awake macaque monkeys using implanted phased-array coils (Janssens et al., 2012) and contrast-agent enhanced fMRI (Vanduffel et al., 2001). This yielded high-resolution (0.6 mm isotropic voxels) maps, close in resolution to most microelectrode retinotopic mapping studies (0.3∼0.5 mm in depth, but 0.8∼1 mm between penetrations parallel to the cortex). Using this technique, we have reliably identified finely-organized, interdigitated V2 stripes in awake behaving macaques, with high intra- and inter-subject reproducibility (Li et al., 2017). Importantly, we calculated fine-grained iso-polar angle and iso-eccentricity contour lines and superimposed these onto the field sign maps, which drastically improved detectability of subtle visual field transitions. Finally, we supplemented the retinotopic maps with population receptive field (pRF) size analyses (Dumoulin and Wandell, 2008), to provide a comprehensive quantitative evaluation of the models. Our results revealed an organization of dorsal third- and fourth-tier visual areas immediately anterior to V2d that largely reconciles macaque and New World monkey models.

## Results

### Three retinotopically distinct sectors in cortex immediately anterior to macaque V2d

Figure 2 displays retinotopic maps acquired from the top half of the left hemisphere of a representative subject (M1) as flattened (A), folded and inflated surfaces (B). Based on field sign reversals, several areal borders can be readily identified. The V1d/V2d border can be recognized as representing the LVM (labeled with red “a” in Figure 2A, left panel) along a clear transition in field sign and a reversal of the polar angle. This border is located along the caudal lip of the lunate sulcus (Figure 2B), which matches its reported anatomical location (see Figure 1A) (Gattass et al., 1981; Lyon and Kaas, 2002c; Roe and Ts’o, 1997). Another LVM (red “b” in Figure 2A, left panel) can be recognized along the prelunate gyrus and forms the posterior V4 border, in agreement with the prevailing V4 macaque model (Figure 1A). Between V2 and V4, both polar angle and eccentricity maps become more distorted and complicated, which has led to the plethora of partitioning schemes described in the introduction. Our data revealed two mirror-image sectors (R1 and R2, colored in blue) immediately rostral to V2d, with a gap (R3, non-mirror image, colored in red) separating them. The gap has been sporadically reported, and has been regarded as a simple interruption of V3d (Gattass et al., 1988; 2015; Kaas et al., 2015), in that scheme corresponding to a combination of R1 and R2. We will argue, however, that the latter two sectors cannot comprise a single visuotopic area and even belong to different hierarchical levels.

**Figure 2.**
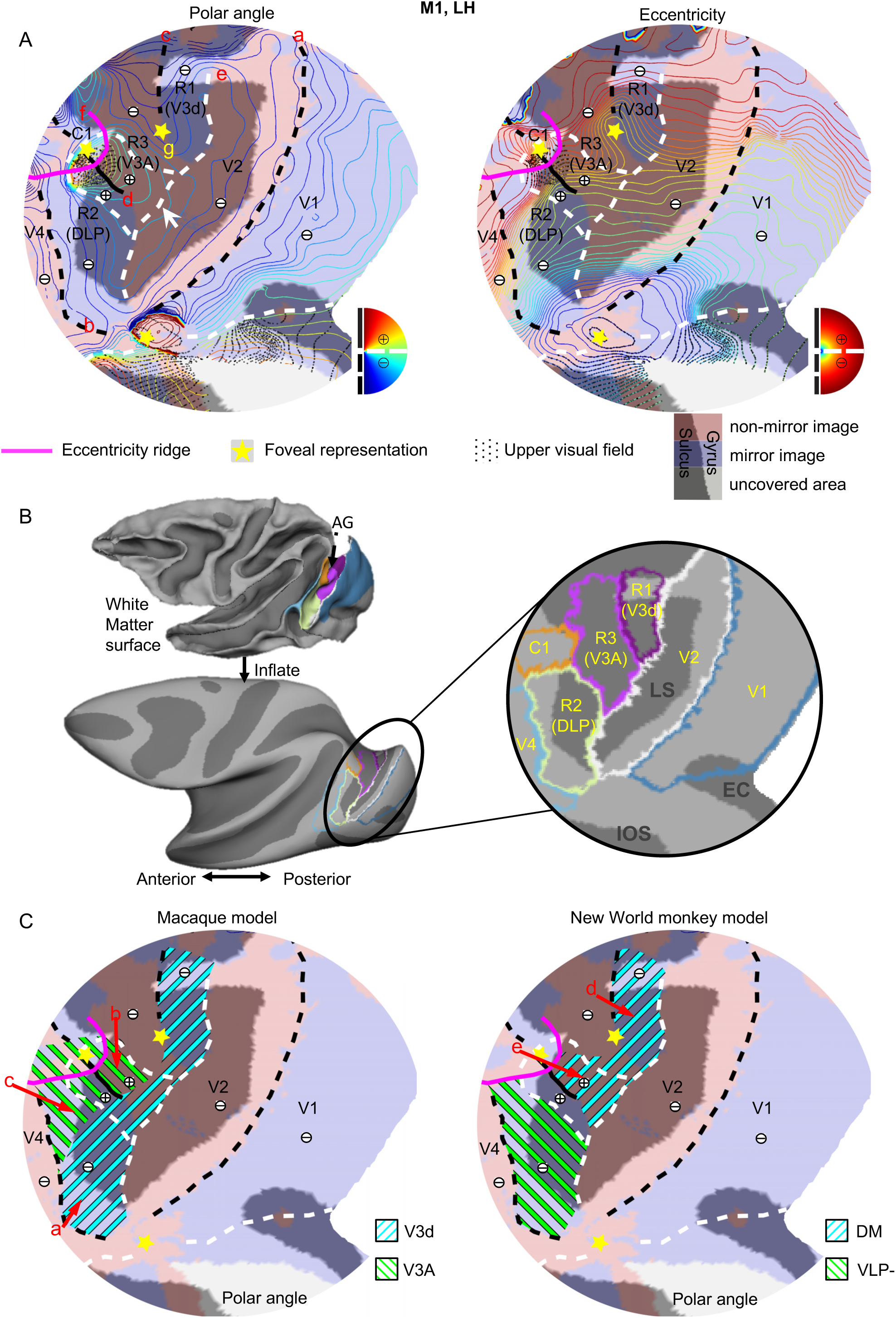
Retinotopic organization of macaque dorsal areas (subject M1). ***A*** Polar angle and eccentricity maps (colored iso-contour lines) are displayed on the field sign map (red and blue shadings) of the left hemisphere (LH) of subject M1. ***B*** Areal outlines are shown on folded, inflated and flat surfaces. ***C*** Reconciling results in ***A*** with existing macaque and New World monkey models leads to improbable areas V3A and DM [areas with different field signs are combined in V3A (b and c) and DM (d and e); the same upper visual field locations are represented twice in V3A (c+ and b)]. LS: lunate sulcus; EC: external calcarine sulcus; IOS: inferior occipital sulcus; AG: annectant gyrus.

#### Sector R1 (putative V3d)

The more dorsomedial region (R1) shares a HM with V2d caudally and a LVM (red “c” in Figure 2A, left panel) with the non-mirror R3-sector, rostrally. Hence R1 represents only the lower contralateral quadrant. This sector is located in the anterior bank of the lunate sulcus near the annectant gyrus (see Figure 2B) and most likely corresponds to the most dorsal portion of V3d as reported in the literature.

#### Sector R2 (putative DLP)

The more ventrolateral region (R2), on the other hand, shares an UVM (red “d” in Figure 2A, left panel) caudally with the non-mirror image region (R3), and a LVM (red “b” in Figure 2A) with another non-mirror image region (V4) rostrally. Hence R2 represents the entire contralateral hemifield. This ventrolateral sector also bends away from V2d, exactly as does VLP in the New World monkey model (Figure 1B). However, R2 is distinct from the marmoset’s VLP because of the UVF representation. Instead, it rather resembles DLP as described in New World Owl monkeys (Figure 1C). Theoretically, the dorsomedial part of this mirror image region (green hatched lines in Figure 2C, red “c”, left panel) could also fit V3A from the macaque model 1 (Figure 1A), as they possess similar visual field representations and cortical locations. However, in that case, the most ventrolateral portion of R2 (red ‘a’ in Figure 2C, left panel) belongs to part of V3d while the UVF representation of R3 (red ‘b’ in Figure 2C, left panel) has to be included in V3A. This requires the unlikely combination of two regions with different field signs in one area (red and blue regions indicated by arrows b and c in Figure 2C, left panel). Moreover, it leads to improbable areal definitions, since the UVF will be represented twice (c and b), which is unlikely within a single retinotopic area. Therefore, a single area DLP as in the New World owl monkeys is the most parsimonious explanation for R2.

#### Sector R3 (putative V3A)

The non-mirror sector R3 contains a representation of the entire contralateral hemifield, with a LVF (R3-) represented dorsomedially at the annectant gyrus, and an UVF (R3+) represented ventrolaterally at the anterior bank of the lunate sulcus, remarkably similar to the V3A originally described by Van Essen and Zeki (1978). It also resembles DM as proposed by Lyon and Kaas (2001; 2002c) in both Old and New World monkeys based on connectivity evidence (Figure 1E). The more dorsomedial part of this sector shares a (near)center of gaze (yellow “g” in Figure 2A, left panel) with R1 (V3d), around which near-foveal iso-eccentricity lines are clearly wrapped. The LVF representation (R3-) resembles VPP-, and the UVF representation (R3+) resembles DI+ as described by Sereno et al. (2015) for owl monkeys, insofar as they represent the same quadrant and have the same non-mirror field sign (Figure 1C). Unlike in owl monkeys, however, we find no evidence for the UVF representations VPP+ and DM+ (Figure 1C) which should appear lateral to the center of gaze. The separation of R3- from R3+ would lead to improbable areas, each representing a quadrant only. Rostral to R3 (putative V3A) and dorsomedial to R2 (DLP), an eccentricity ridge (red “f” in Figure 2A, left panel) can be recognized around another foveal representation (near C1 in Figure 2A, left panel). This ridge aligns well with the transition of visual field sign at the dorsomedial border of R2 and the transition at the rostral border of R3 (V3A), thereby separating a small cluster of areas (C1) from both R2 and R3. In most hemispheres (Figure 3 and 4), this C1 cluster contains a double representation of the entire contralateral hemifield, and hence consists of more than one visual area. C1 will not be discussed further, except to say that it appears similar to PP in owl monkeys (Sereno et al., 2015) (Figure 1C). C1 might also overlap with a histologically-defined intermediate area (IA) because of their nearly identical locations (Van Der Gucht et al., 2006). Indeed, both areas are located dorsal to V4d but ventral to the dorsal prelunate area (DP) on the prelunate gyrus.

**Figure 3.**
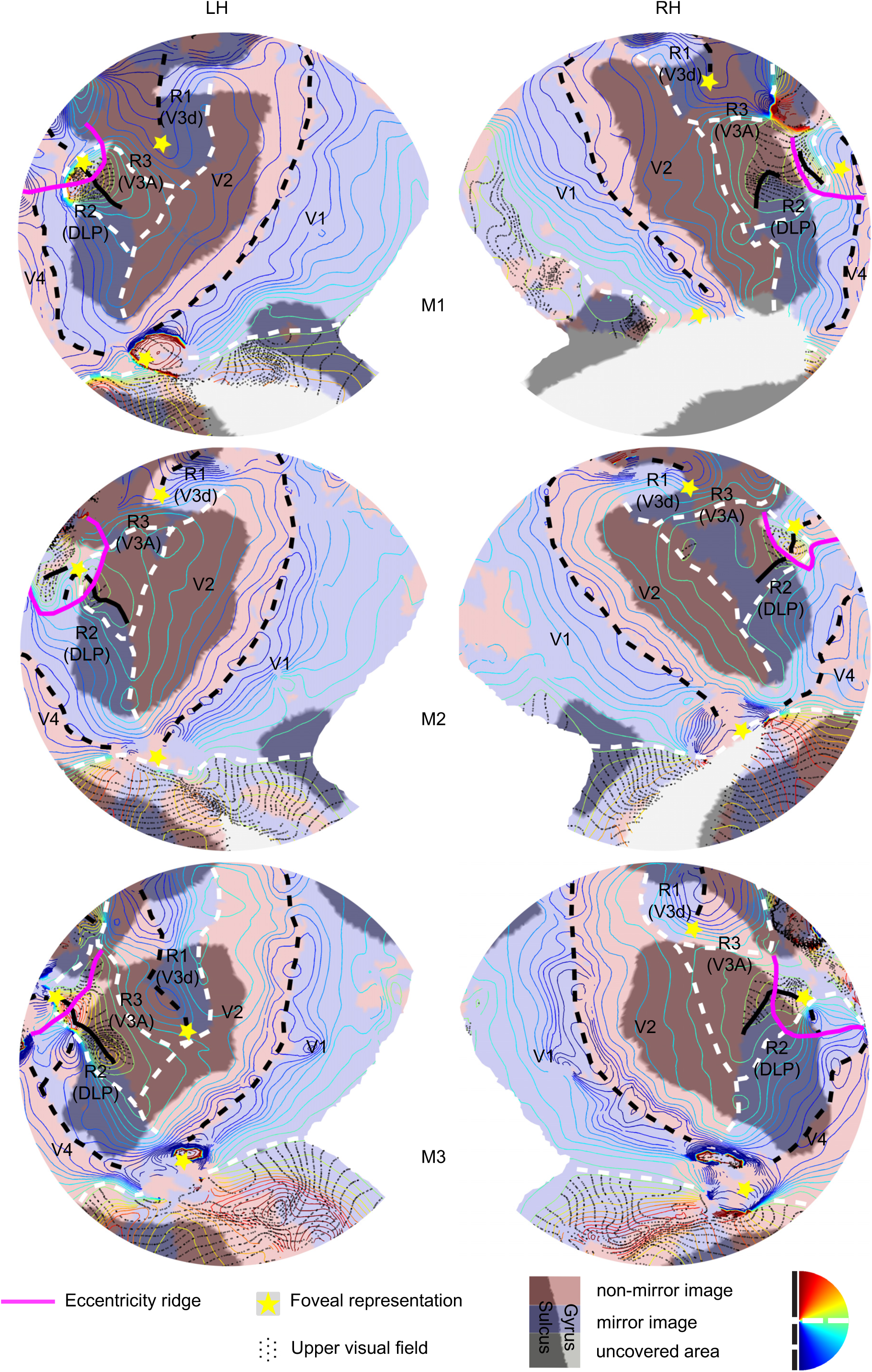
Polar angle organization of the dorsal areas in 6 hemispheres (M1-3). LH, left hemisphere; RH, right hemisphere.

**Figure 4.**
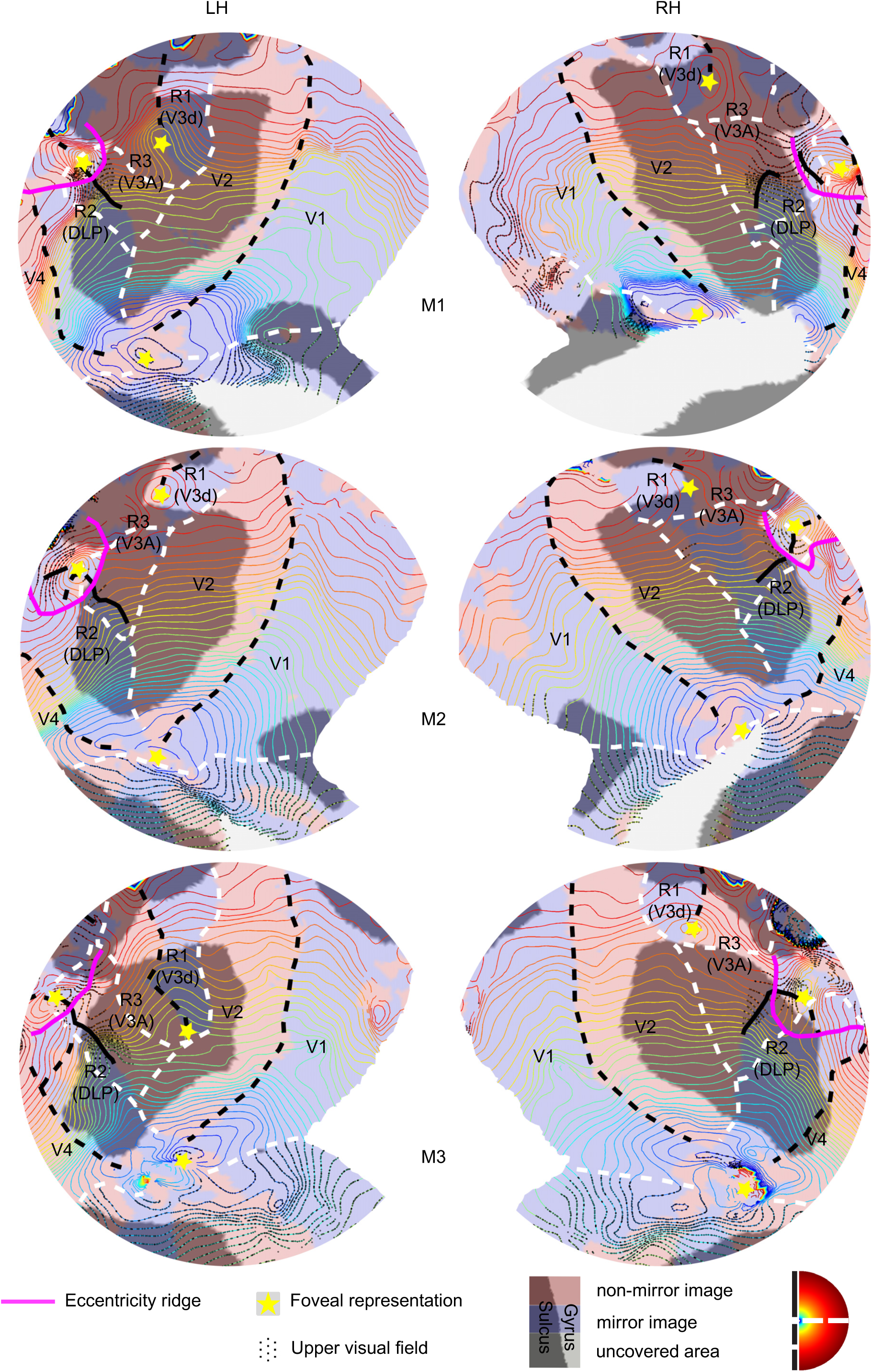
Eccentricity organization of the dorsal areas in 6 hemispheres (M1-3). LH, left hemisphere; RH, right hemisphere.

#### Consistent retinotopic layout of dorsal visual area across 6 hemispheres

The topographic organization described in detail for the left hemisphere of M1 is surprisingly consistent across all 6 hemispheres, as displayed in Figures 3 and 4. Crucial features include the non-mirror image gap (R3) between R1 and R2 and an UVF representation that covers both the mirror (R2) and non-mirror (R3) sectors indicating that there are two, rather than one, upper-field quadrants. Only the right hemisphere of subject M2 shows no pronounced gap. In general, our results point to an organization for dorsal visual cortex differing from macaque models 1 and 3, as presented in Figure 1. A ventrolateral mirror image region R2, originally assigned to V3d and V3A in the macaque model 1, fits best with area DLP of owl monkeys. Furthermore, V3d (R1) extends only along the most dorsomedial portion of the anterior V2d border. R3, possessing a full hemifield representation, is wedged in between R1 and R2 and most likely corresponds to V3A. This organization resembles that seen in New World monkeys (Figure 1B, C), except that, different quadrants are combined to form areas with a complete contralateral visual field representation. Below, we will use quantitative pRF size measurements to corroborate our hypothesis that R2 and R1 do not belong to the same area, and should even be assigned to different hierarchical levels.

### pRF sizes in R2 (putative DLP) and R1 (putative V3d)

To objectively evaluate the various models, we estimated pRF sizes within each voxel (Dumoulin and Wandell, 2008). More specifically, we tested whether R2 and R1 belong to V3d as predicted by the classical macaque models 1 and 3. Alternatively, they consist of different areas (i.e. DLP and V3d, respectively), as suggested above based on our high-resolution retinotopic maps. Figure 5 displays individual (panel A) and average pRF sizes (B) as a function of eccentricity, for each sector of interest, and including V2d. In all areas, the pRF size increases with eccentricity, and across all individual subjects, the pRF size at higher eccentricities increases from caudal (e.g. V2d) to rostral areas (e.g. V4d). Furthermore, pRF size at high eccentricities of R2 is much larger than that of R1 in each subject.

**Figure 5.**
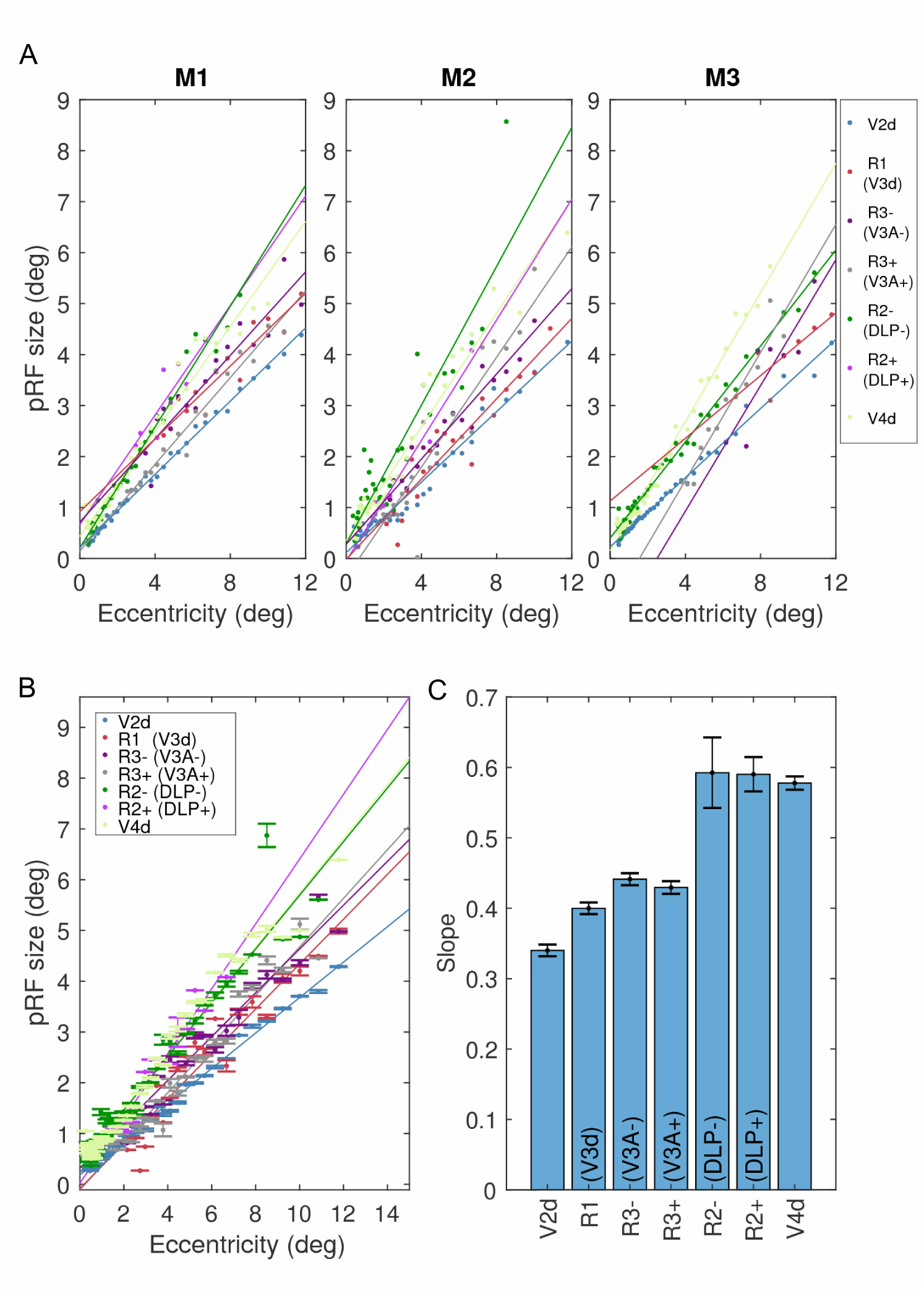
pRF sizes of all dorsal visual areas discussed in the present study. ***A-B*** Individual (***A***) and average pRF sizes (***B***) as function of eccentricity, in each sector of interest rostral to and including V2d. ***C*** Slope estimates (mean and standard error) from the best-fitted linear mixed-effect model.

The linear relationship between eccentricity and receptive field size allows the use of a linear mixed-effect analysis to statistically compare pRF sizes across areas. This analysis showed that a model with a common intercept but separate slope terms best describes the data [Bayesian information criterion (BIC) = 479.09], compared to a simpler model without the slope terms in the random effects (BIC = 497.53), or a more complex model varying the common intercept term in the fixed effect as separate intercept terms for different areas (BIC = 495.09). Hence, slopes can be compared across areas to quantify areal differences based on pRF size. The best model provided an excellent fit of the data, explaining 94% of the variance (Adjusted r-squared = 0.941).

Figure 5C displays the model’s slope estimates of all dorsal visual areas discussed in the present study. A pairwise comparison of the slopes showed that pRF size is smallest in V2d and largest in V4d (V2d < V3d < V4d, *p*s<10^-4^, FDR corrected), consistent with most previous findings (Boussaoud et al., 1991; Desimone and Schein, 1987; Gattass et al., 1981; 1988; Hubel et al., 2015; Rosa and Tweedale, 2000a). There is no difference between the slopes of the two quadrants of R3 (V3A) (p = 0.34, FDR corrected). Importantly, the slope of R3+ is significantly smaller compared to R2+ (p < 0.005, FDR corrected), adding to the field-sign based evidence that these are 2 independent upper field quadrants. Moreover, R3’s slopes are significantly larger than those of V2d and R1 (V3d) (*p*s < 0.02, FDR corrected), but smaller than V4d (*p*s < 10^-4^, FDR corrected). The slope of the two R2 quadrants are indistinguishable (p = 0.968, FDR corrected). Critically, they are significantly larger than those of R1 (V3d) (p < 10^-3^, FDR corrected), suggesting that R2 corresponds to another area rather than being part of V3d. Interestingly, R2’s slopes are equal to that of V4d (*p*s > 0.6, FDR corrected), suggesting areas at the same hierarchical level (4^th^-tier) of the visual system.

According to the macaque models 1 and 3, only the more ventrolateral part of R2 should be considered as part of V3d (blue hatched area ‘a’ in Figure 2C, left panel), whereas the more dorsomedial part would belong to V3A (green hatched area ‘c’ in Figure 2C, left panel). Therefore, we performed an additional analysis, dividing R2 into two parts, and comparing their respective pRF sizes with R1. The macaque models 1 and 3 predict pRF sizes for the ventrolateral portion of R2 similar to those of R1. If true, and based on the pRF size of the entire extent of R2 [green (c) and blue (a) hatched areas in Figure 2C], its dorsomedial portion should have much larger pRF sizes compared to its ventrolateral portion. However, this is not the case. Figure 6A displays the pRF size in R2 (DLP) and R1 (V3d) as a function of eccentricity, with V2d as a reference. The vertical dashed line represents 2.5° eccentricity, which corresponds roughly to the eccentricity separating the LVF representation of V3d from V3A in macaque model 1 (Gattass et al., 1988), and DM from VLP in the New World monkey model (Rosa and Tweedale, 2000a; 2005). Unlike predictions based on macaque models 1 and 3, pRF sizes in R2 (DLP) on opposite sides of this eccentricity line show similar linear trends for low and high eccentricities, like those observed in V2d. Furthermore, even the slope of the foveal portion of R2 (DLP) seems large compared to that of R1 (V3d).

**Figure 6.**
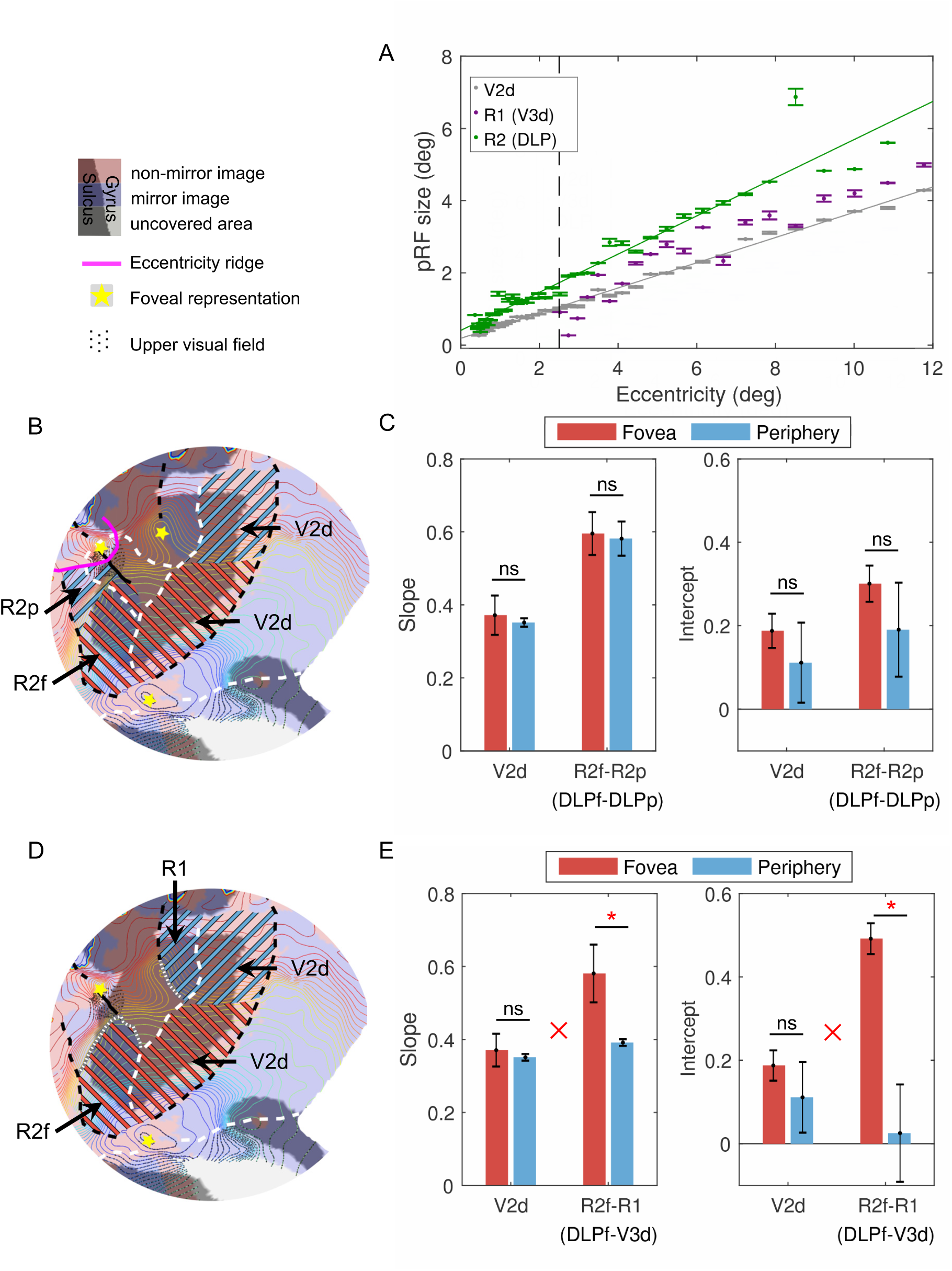
Different pRF sizes in R1 (V3d) and R2 (DLP). ***A*** pRF sizes as function of eccentricity in V2d, R1 (V3d) and R2 (DLP). The vertical dashed line represents 2.5° eccentricity, corresponding roughly to the eccentricity separating the LVF representation of V3d from V3A in macaque model 1 (Gattass et al., 1988; 2015), and DM from VLP in the New World monkey model (Rosa and Tweedale, 2000a; 2005). The pRF size in R2 (DLP) separated by this eccentricity line shows a similar linear trend with eccentricity, as in V2d. ***B-C*** Slope and intercept estimates in foveal R2 (R2f/DLPf) and V2d (red hatching in ***B***) compared with their peripheral counterparts (R2p/DLPp and V2d) (blue hatching in ***B***). ***D-E*** Slope and intercept estimates in foveal R2 (R2f/DLPf) and V2d (red hatching in ***D***) compared with R1 (V3d) and peripheral V2d (blue hatching in ***D***). *, significant pairwise comparison (p < 0.05); ns, no significant pairwise comparison (p > 0.05); red “×”, significant two-way interaction (p < 0.05).

In Figures 6B and 6C, the pRF sizes in the foveal (R2f/DLPf, eccentricity < 2.5°) and peripheral portions (R2p/DLPp, eccentricity > 2.5°) of R2 (DLP) were statistically compared using two linear mixed-effect models, using V2d as a reference. In the first model, a common intercept but different slope terms were applied to the foveal and peripheral representations of each area, and the resulting slope estimates were compared. In the second model, the slope but not the intercept terms were held equal, and intercept estimates were compared. The results of both models showed highly similar results (*p*s > 0.2) for the foveal and peripheral portions of R2 (DLP), comparable to reference area V2d. Furthermore, differences in slopes or intercepts between foveal and peripheral representations are indistinguishable for R2 (DLP) and V2d (2-way interaction, *p*s > 0.7). In Figure 6D and 6E, a similar comparison was made between R1 (V3d) and the foveal portions of R2 (R2f/DLPf). In this case, both the slope and intercept estimates of R2f (DLPf) are significantly greater than in R1 (V3d) (*p*s < 0.02) and these differences significantly exceed those between foveal and peripheral V2d (2-way interaction: slope estimates, p = 0.0477; intercept estimates, p = 0.0035). Moreover, the same distinction between R2f (DLPf) and R1 (V3d) is obtained when much smaller eccentricity values are taken as the cutoff (e.g. 1°, 2-way interaction between intercept estimates, p = 0.0072). Hence, these findings cannot be attributed to a larger area being assigned to the foveal portion of R2 (DLP) versus R1 (V3d) by using a 2.5° eccentricity cutoff. These data add to the evidence that R2 (DLP) is quantitatively different from R1 (V3d), and strongly argue against an elongated V3d, as proposed in macaque models 1 and 3 (Figure 1A, E).

### Does R2 (DLP) fit the New World monkey model for third-tier areas?

To test whether the LVF quadrant of R2 (R2-/DLP-) may be considered a continuation of V3v as in a New World monkey model (Figure 1B), we also acquired retinotopic maps for ventral areas in all subjects and compared pRF sizes in R2- (DLP-) and V3v. Figure 7A displays retinotopic maps of ventral areas in subject M1 acquired in separate sessions during which only the bottom half of the brain was covered due to limited slice coverage. Compared to dorsal areas, the retinotopic organization of ventral areas is less complex, and areal boundaries can be straightforwardly delineated along field sign reversals. Specifically, rostral to V2v in each hemisphere, a single narrow band of a mirror image representation defines area V3v, in which the visual field moves from the HM (posterior) to the UVM (anterior). Rostral to V3v, the visual field representation moves from the UVM back to the HM, mirroring that of V3v. These data exactly fit the retinotopic organization of V4v described in the literature (Boussaoud et al., 1991; Gattass et al., 1988; Newsome et al., 1986; Piñon et al., 1998). Figures 7B-C display individual and average pRF sizes, respectively, of these ventral areas together with their dorsal counterparts. pRF sizes increase linearly with eccentricity in all tested areas. A linear mixed-effect model, with fixed and random effects including both common-intercept and separate-slope terms for each area, best fitted the data (BIC = 402.87), compared to the simpler (BIC = 464.89) and more complicated models (BIC = 433.76). The slope estimates (Figure 7D) from the best model indicated similar receptive field sizes in ventral and dorsal V2 and V4, respectively (*p*s > 0.6, FDR corrected), highly consistent with previous single-cell recording studies (Gattass et al., 1981; 1988; Piñon et al., 1998; Rosa et al., 1997). Similarly, the slope of V3v equals that of R1 (V3d) (p = 0.144, FDR corrected), but is significantly smaller than that of R2- (DLP-) (p < 0.001, FDR corrected). This suggests that R2 (DLP) is a higher level area relative to V3v. Hence, the latter two areas cannot be combined to form a monkey equivalent of VLP as proposed in the New World marmoset model (Figure 1B).

**Figure 7.**
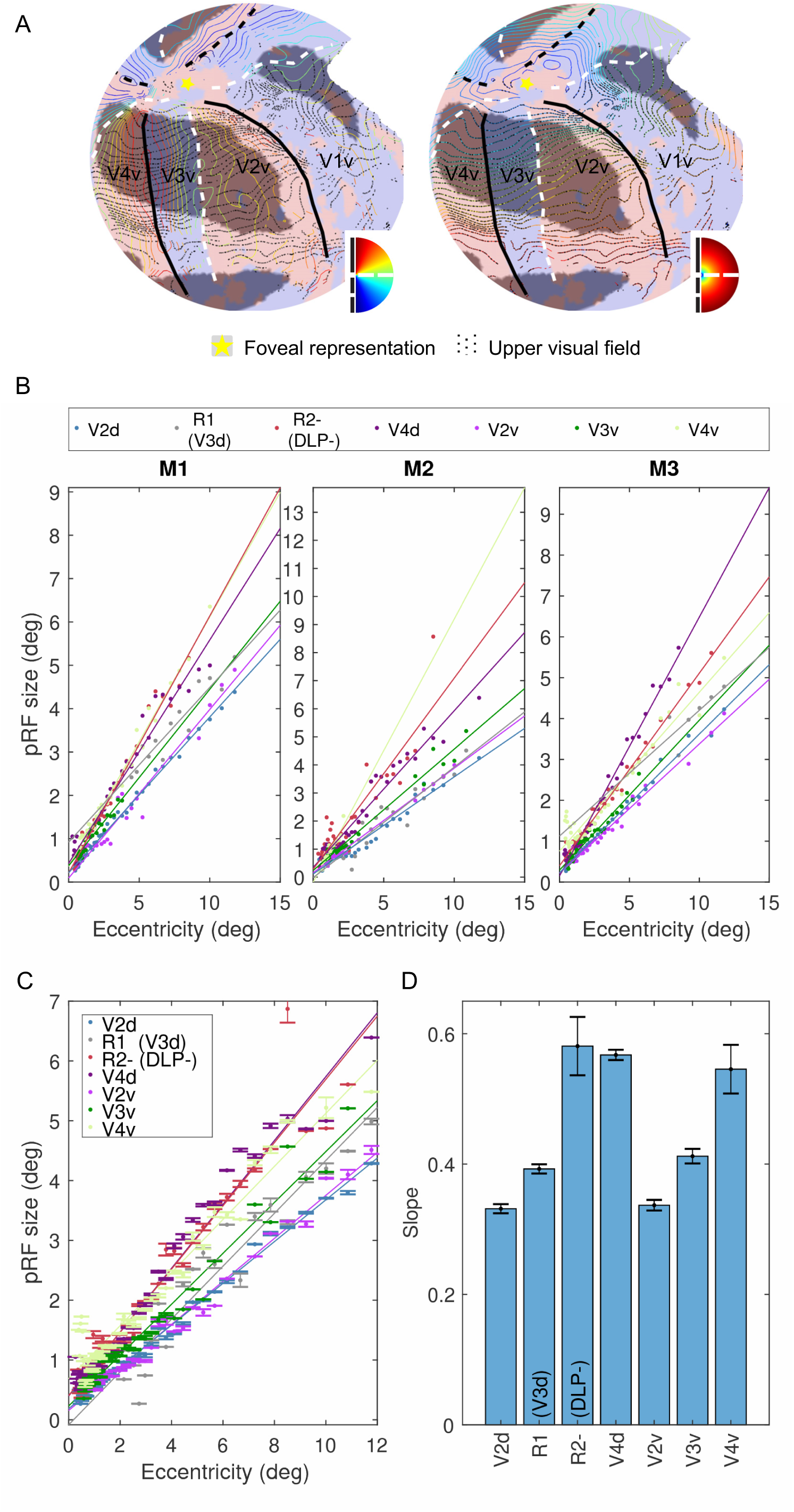
Different pRF sizes in V3v and R2 (DLP). ***A*** Retinotopic organization of the ventral areas of the left hemisphere of subject M1. ***B*** Individual pRF sizes of each quadrant in each monkey. ***C*** Average pRF sizes and slope estimates between eccentricity and pRF size in different areas. pRF size differences: V2d = V2v < V3d = V3v < DLP = V4d = V4v; =, p>0.05; <, p < 0.05, FDR corrected.

## Discussion

Our sub-mm retinotopic mapping of monkey cortex demonstrated multiple retinotopically-organized areas immediately rostral to V2d. The most dorsomedial area (R1) contains a mirrored visual field representation of V2d and its receptive field size equals to that of V3v, exactly as previously described for V3d. Hence, we consider it as the only sector that can correspond to V3d proper. Rostral to the middle sector of V2d, a non-mirror image area (R3) is located at the annectant gyrus, with a LVM and UVM at its posterior and anterior borders, respectively. Hence, the organization of R3 is surprisingly similar to V3A as initially described by Van Essen and Zeki (1978). The most lateral area (R2) curves away from V2d and contains a representation of the entire contralateral hemifield, exactly fitting DLP, as described for New World owl monkeys (Sereno et al., 2015). Its foveal portion directly abuts V2d and is separated from V3d by a gap. Considering R1 and R2 together, they resemble a discontinuous V3d as proposed by macaque models 1 and 3. One might be tempted to argue that its LVF representation resembles dorsal VLP, as proposed in the New World marmoset model. However, R2’s pRF size is significantly larger than that of V3d (R1) and V3v. Hence R2 can be considered neither a part of V3d, nor a LVF continuation of V3v as proposed by the New World marmoset model. Receptive field size in both its visual quadrants, however, is significantly larger than V3A, but quite similar to that of V4, suggesting that R2 (DLP) is a fourth-, rather than third-tier area. Together, the results revealed a topographic organization for macaque dorsal visual areas rostral to V2d that substantially differed from the most widely accepted models, including V3d, V3A, and V4d.

### V3d

Our results confirm the existence of a V3d (=R1) in macaque monkeys. A New World monkey DM would require a combination of R1 (V3d) with the UVF representation of R3 (V3A+) (see Figure 2C, right panel). However, R1 (V3d) and R3+ (V3A+) have different visual field representations (mirror vs. non-mirror image) and different receptive field sizes, and thus are unlikely parts of the same area. According to the summary maps in Figure 2 of Rosa et al. (2009) and Figure 1b of Jeffs (2013), the progression of eccentricity is the same in both DM- and DM+, whereas the polar angle progression is reversed along a caudo-rostral axis (from the HM to the LVM in DM- but to the UVM in DM+). Hence, the visual field representation is also different in DM- and DM+ (a mirror and non-mirror image representation in DM- and DM+, respectively) in the New World marmoset monkey model, exactly as in V3d and V3A+. The DM in the model of Kaas and co-workers (Kaas et al., 2015; Stepniewska et al., 2005) (Figure 1E), however, is more similar to R3 (V3A). Therefore, they proposed a V3d lying between V2d and DM, which corresponds exactly to R1 (V3d) as in our results.

A putative New World marmoset model also requires merging R2- (DLP-) with V3v, since V3v alone would then constitute an improbable area in which only one quadrant is represented (Angelucci and Rosa, 2015; Rosa and Tweedale, 2000a; 2005). R2 (DLP) contains the same mirror image representation of the visual field as V3v, however, its receptive field size is significantly larger than V3v. This result, although seemingly in conflict with the marmoset model having a similar receptive field size in ventral and dorsal VLP (Rosa and Tweedale, 2000a), is quite consistent with results in macaques describing a posteromedial area (PM) (Maguire and Baizer, 1984). This area resembles R2- (DLP-), and contains neurons with receptive fields similar in size to V4 neurons. Thus, in macaque monkeys, converging evidence suggests that R2 (DLP) is a higher-level area, distinct from V3v. R2 therefore cannot be combined with V3v, as proposed for New World marmoset monkeys. This leaves V3d, with the same mirror image representation and pRF size as V3v, as the only quadrant representation that can be combined with V3v to form a complete area V3.

However, unlike macaque models 1 and 3, in which V3d extends along most of the V2d border, our V3d (R1) is reduced in size and borders V2d only anteromedially. The foveal part of V3d from macaque models 1 and 3 belongs to another area, i.e. DLP (R2). Since this portion of the lunate sulcus represents the fovea, has small receptive field sizes, and contains the same visual field progression as V3d, it is unsurprising that foveal R2 has previously been considered part of V3d (Gattass et al., 1988).

One could argue that a pinched portion of V3d, reduced to 50% of its width compared to R1, could still exist between R3+ and neighboring V2. However, based on our simulations (Figure S2), we would have been able to resolve this thin sliver of V3d, given our sampling resolution and an SNR of 40 (Figure S2A). In fact, V3d needs to be narrower than 0.3 mm to remain undetected by our field sign analysis (Figure S2B). Therefore, it is unlikely that we missed a portion of V3d, as previously depicted in the macaque model, a consequence of coarse sampling and smoothness of fMRI data.

The observed organization of V3d, although a departure from the traditional macaque model, is in fact largely compatible with connectivity data. First, it fits quite well with connectivity evidence from Lyon and Kaas (2002c) confirming the existence of V3d in macaque monkeys, since all tracer injections were made at eccentricity representations > 5°. In fact, this also holds true for the majority of injections in other tracing studies [e.g. Gattass et al. (1997) and Ungerleider et al. (2008)].

Secondly, the new model explains connectivity patterns that cannot be accounted for by the old macaque models (1 and 3). Stepniewska and Kaas (1996) injected a series of tracers in macaque V2 and found a connectivity pattern in dorsal areas surprisingly similar to that of New World monkeys. The organization of their lower field DM (DM-) is similar to our V3d, in the sense that its ventral extension ends at the anterior V2 border near 2° eccentricity, and in that a similar center of gaze is represented at its ventral tip immediately rostral to V2d, as corroborated by our results. This result is difficult to reconcile by macaque models 1 and 3, since the same foveal visual field would be duplicated in the same area.

Finally, in a connectivity study by Ungerleider et al. (2008), a series of anterograde and retrograde tracers were injected at different eccentricities in V4. This yielded a topographically organized connectivity pattern in V3d, with a clear separation between foveal and more peripheral projections from V4 to V3d, thus fitting our model instead of the classical macaque model. Furthermore, there is a clear difference in laminar distribution of labelled cells and terminals in V3d after foveal versus peripheral V4 injections, with the latter projections consistently classified as “feedback”, whereas those identified after foveal injections being of the “intermediate” type (see their table 2). This distinction cannot be explained by macaque models 1 and 3, since a homogeneous distribution of laminar projection patterns should be expected in a single area. However, the connectivity results are entirely consistent with the present results, whereby DLP and V3d are different areas separated by a clear gap. Thus, it is very likely that the foveal portion of DLP has previously been misassigned to area V3d.

### V3A

Rostral to V3d, we observed another third-tier area, V3A (R3), highly similar to the initial description given by Van Essen and Zeki (1978). This V3A occupies most of the annectant gyrus, sharing a LVM with V3d caudomedially and an UVM with DLP (R2) rostrolaterally. The pRF size of this V3A is slightly larger than that of V3, consistent with the observations of Van Essen and Zeki (1978). However, unlike the rather complex retinotopic organization described by Van Essen and Zeki (1978), we observed a single, slightly warped but orderly representation of the entire contralateral hemifield in this area, whereby the receptive fields move from the lower to the upper quadrant along the caudo-rostral axis.

The most widely-accepted macaque model (model 1, Figure 1A), however, describes a different retinotopic organization for V3A, whereby the UVF representation borders V3d posteriorly, near the narrow V3d zone, whereas the LVF representation borders V4 anteriorly. This arrangement is significantly different from our observations and those of Van Essen and Zeki (1978), but may be explained if one combines peripheral DLP with the UVF representation of our V3A (R3+) (Figure 2C, left panel). Exactly as with the V3A of Gattass et al. (1988), this combination of quarter fields is located mainly at the anterior bank of the lunate sulcus, largely avoiding the annectant gyrus. Moreover, we observed significant differences in field sign and pRF size between R2+ and R3+, indicating that they belong to separate areas.

The retinotopic organization in V3A as described by Gattass et al. (1988) was reported in two studies of Nakamura and Colby (2000; 2002) when recordings were made in the annectant gyrus. However, these results may have been misinterpreted due to the complex folding of the annectant gyrus. According to the folded surface of a representative subject displayed in our Figure 2B, progression of recording sites along the annectant gyrus on sagittal sections as sites 12 to 17 [supporting Figure 9A of Nakamura and Colby (2002)] can result in a progression path runs from lateral to medial on the flat surface (see yellow arrowhead in Figure S3), which, according to our results, will produce a progression from the horizontal to the vertical meridian. This is very similar to the visual field shifts shown in supporting Figure 9B of Nakamura and Colby (2002), suggesting that the retinotopic organization of V3A observed in their study is more consistent with our results, compared to model 1. In two other single-cell recording studies, Galletti and co-workers (Galletti and Battaglini, 1989; Galletti et al., 1990) also described a retinotopic organization in V3A resembling our results and those of Van Essen and Zeki (1978). In summary, our results confirm the existence of V3A as originally defined by Van Essen and Zeki (1978) and several following studies (Galletti and Battaglini, 1989; Galletti et al., 1990; Nakamura and Colby, 2000; 2002; Zeki, 1978a). However, it is different from V3A described in model 1 (Gattass et al., 1988; 2015).

### V4

Anterior to DLP and V3v, we observed a non-mirror image area extending dorsally from the prelunate gyrus to the posterior bank of the STS and ventrally from the anterior bank of inferior occipital sulcus onto the inferior occipital gyrus. This area shares a VM and a HM with posterior and anterior areas, respectively, very similar to area V4 as described in macaque model 1 (Gattass et al., 1988; 2015). Since V4 has been widely accepted in this form, we followed this designation here. It should be emphasized, however, that in its original description by Zeki (1978b), “V4” referred to a “fourth visual complex” including multiple areas: one located in the anterior bank of the lunate sulcus and another extending from the anterior bank of the lunate sulcus onto the prelunate gyrus. The location of the former area is very similar to our DLP, and the latter is similar to the dorsal portion of our V4 (V4d). These two areas were grouped, since the properties of the cells in these two areas were very similar (Zeki, 1978b). Consistently, we observed similar pRF sizes in DLP and V4. The combination of V4d and DLP conforms to the same retinotopic organization as observed for the V4-complex by Van Essen and Zeki (1978), since they described an UVM representation at the V3A/V4-complex border, and that the same visual field is represented twice in the V4-complex. Therefore, we suggest that DLP and V4 of the present study are parts of Zeki’s V4-complex.

Our results concerning the dorsal fourth-tier areas are also consistent with Baizer and Maguire (1983), Maguire and Baizer (1984), and Youakim et al. (2001), who described two areas, PM and AL (anterolateral area), occupying the macaque prelunate gyrus. These are separated by a LVM and their neurons have the same receptive field size, exactly as we found in area DLP and V4d. The main difference is the upper visual quadrant representation in area DLP which is lacking in PM, although they claim it is present just outside PM’s border (Maguire and Baizer, 1984).

Our observation of two dorsal fourth-tier areas around the prelunate gyrus is also consistent with the connectivity results observed by Stepniewska and Kaas (1996) and Stepniewska et al. (2005) in macaques. They described two distinct projection fields from the anterior bank of the lunate sulcus onto the prelunate gyrus after tracer injections in V2. These two projection fields are similar to the V4 and V4A observed by Zeki (1971) after lesions made in V2 and V3, but were designated DLr and DLc following the terminology used in New World monkeys.

These results, however, are inconsistent with connectivity results observed by Gattass et al. (1997), who observed a single projection field in their V4d after injections of tracers in V2d (except in their case 8). Different definitions of area V3 and V3A have likely caused discrepant interpretations of these studies. According to our results, area V3A and the foveal portion of V3 in the macaque model of Gattass et al. (1988; 2015) correspond to our DLP or the dorsal portion of DLr of Stepniewska and Kaas (1996) and Stepniewska et al. (2005), whereas Gattass’ area V4d corresponds to our V4d or the dorsal portion of DLc of Stepniewska and Kaas (1996). Therefore, it is unsurprising that Gattass et al. observed only a single projection field in their V4d, as they assigned the other projection field to other areas (e.g. V3A).

Our description of the ventral fourth-tier area, however, differs from that proposed by Stepniewska and Kaas (1996) and Stepniewska et al. (2005), as we observed only a ventral counterpart for V4d (quarter field) and obviously not for DLP (which is already a hemifield), while Stepniewska and Kaas described upper field quadrants for both DLr and DLc (Figure 1E). Zeki (1971) also described two projection fields from V2v and V3v immediately rostral to V3v, i.e. V4 and V4A. These regions correspond to our V4v and another mirror image area rostral to V4v, which we have also named V4A in our previous studies (Janssens et al., 2014; Kolster et al., 2014) (see also Arcaro and Livingstone, 2017a; 2017b; Pigarev et al., 2002). These two areas are very similar to the ventral portions of DLr and DLc proposed by Stepniewska and Kaas (1996) and Stepniewska et al. (2005) since they have similar connectivity profiles with V2 and similar cortical locations relative to other areas. However, V4v and V4A cannot be considered as the ventral counterparts of DLP and V4d respectively since their visual fields are entirely different (i.e. mirror versus non-mirror image representations). It is possible that these areas are incorrectly combined by Stepniewska and Kaas (1996), Stepniewska et al. (2005) and Zeki (1971), since their detailed visual topography (especially polar angle representation) is difficult to discern based on connectivity patterns only. Furthermore, Zeki (1978b) grouped V4A and V4 together based on the spike data from their dorsal portions, which in fact correspond to our DLP and V4. Whether ventral V4A can be considered as another fourth-tier area is still an open question. According to our previous results (Kolster et al., 2014), this area has larger pRF sizes than V4. But further testing is required to confirm that V4A is indeed a higher-tier area.

### Comparison Old and New World monkeys

A direct comparison of our results (Figure 8A) with the most recent maps of New World marmoset and owl monkeys (Figure 8B-C) revealed remarkably similar polar angle and eccentricity representations in dorsal visual cortex. Within the 12° of eccentricity that we covered, two HM representations are connected perpendicularly with the HM constituting the rostral V2d border in each species. These HM’s separate visual cortex immediately rostral to V2d into three zones: a medial (red outline) and lateral (blue outline) LVF separated by an UVF representation (purple outline). The medial LVF zone is characterized by a LVM in each species, splitting V3d- from V3A- in macaques, DM- from VPP- in owl monkeys and DM- from DA- in marmosets. The UVF zone has a complicated visual field organization, leading to distinct partitioning schemes in different species. However, an UVM is observed in each species separating two sets of upper quadrant representations (i.e. DLP+ in macaque and owl monkeys and DI+ in marmosets, separated from V3A+ in macaques, DI+, DM+ and VPP+ in owl monkeys and DM+ and DA+ in marmosets). The most lateral LVF zone contains a double representation of the contralateral lower visual quadrant in all species. A LVM represents the anterior border of the posterior lower quadrant (DLP- in macaque and owl monkeys, and VLP- in marmosets) shared with another fourth-tier area located more rostrally (V4d in macaques, and V4-like areas DLi and VLA in owl monkeys and marmosets, respectively). Besides the meridians, foveal representations can be retrieved at highly similar topographic locations relative to the medial HM in each species. Hence, the general visual topographic organization of dorsal visual cortex exhibits surprisingly more similarity in Old and New World monkeys than was previously appreciated.

**Figure 8.**
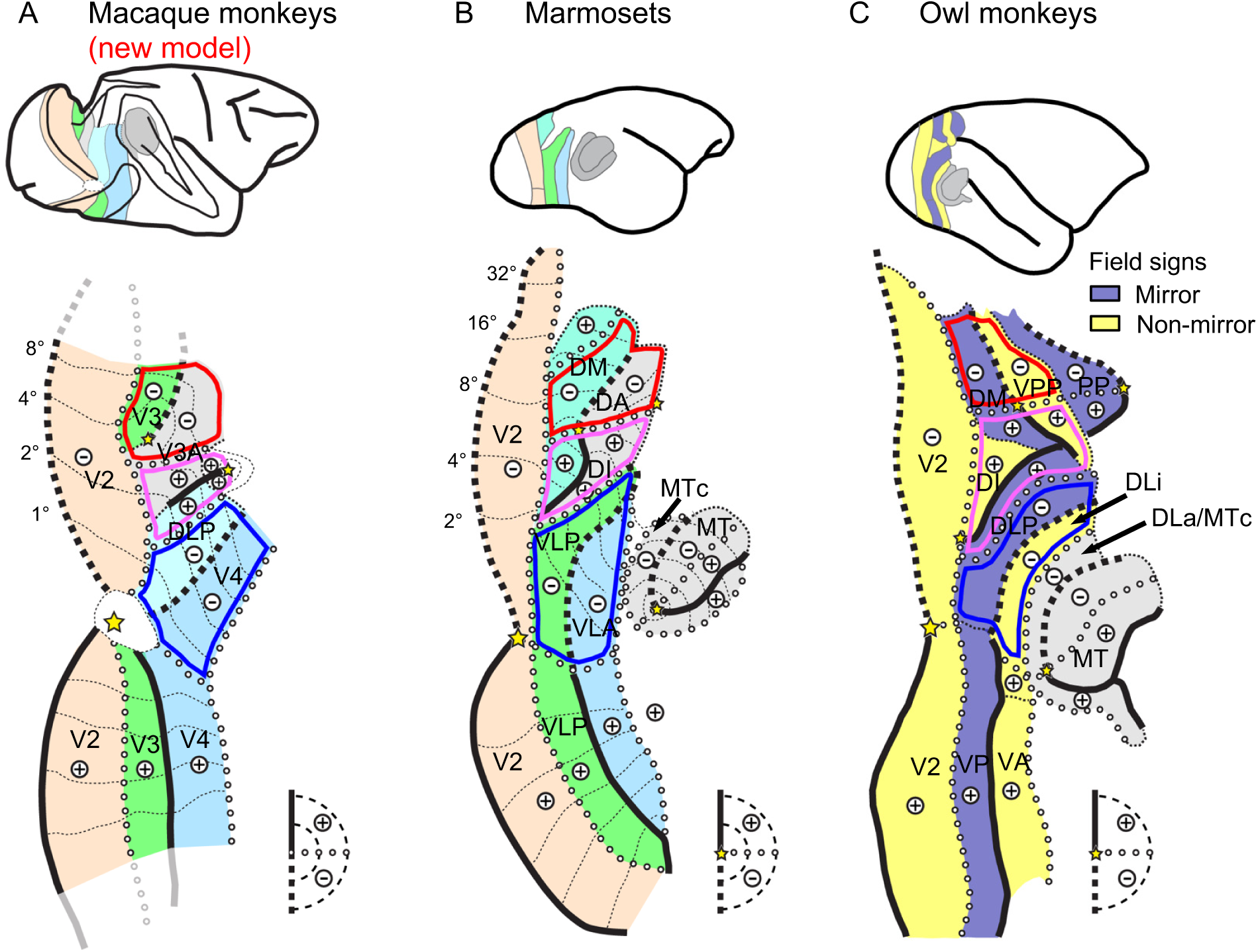
Comparison Old and New World monkeys. A direct comparison of our results (***A***) with the most recent maps of New World marmoset [adapted from Figure 2 of Rosa and Tweedale (2005)] (***B***) and owl monkeys [adapted from Figure 2 of Sereno et al. (2015)] (***C***) suggests that the retinotopic organization of cortex rostral to V2d is evolutionarily largely conserved in Old and New World monkeys.

## Conclusion

Our results reveal a new retinotopic model of the dorsal third- and fourth-tier areas in macaque monkeys. In this model, V3d is reduced in size and borders V2d only anteromedially, similar to DM- in New World monkeys. Area V3A occupies most of the annectant gyrus and shares a LVM with V3 caudomedially and an UVM with a fourth-tier area DLP ventrolaterally. This is exactly as initially proposed by Van Essen and Zeki (1978), but is unlike the macaque model shown in Figure 1A, with an UVF representation posteriorly and a LVM shared with area V4d. Area DLP, an area considered either a combination of V3d and V3A, or a continuation of area V3v, constitutes a fourth-tier area bordering V4d. The overall visuotopic organization of these areas is remarkably similar in Old and New World monkeys, suggesting that this organization is evolutionarily preserved. Future high-resolution imaging is required to ascertain whether a similar organization may exist in humans, as was implied, but not discussed, in the very first phase-encoded retinotopic mapping study of Sereno et al. (1995) (their figure 3).

## Acknowledgments

The authors thank A. Coeman, C. Fransen, P. Kayenbergh, I. Puttemans, S. De Pril, A. Hermans, G. Meulemans, W. Depuydt, C. Ulens, S. Verstraeten, and M. Depaep for technical and administrative support. The authors also thank S. Raiguel for his comments on the manuscript. This work was supported by the Research Foundation Flanders (FWO-Flanders) G0B8617N, G0D5817N, G0A5613N and Odysseus G0007.12; KU Leuven Programme Financing PFV/10/008 and C14/17/109; European Union’s Horizon 2020 Framework Programme for Research and Innovation under Grant Agreement No 720270 (Human Brain Project SGA1), and the Hercules foundation. Q.Z. is a postdoctoral fellow of the FWO-Flanders.

## Author contributions

W.V and Q.Z designed the experiments. W.V performed the surgeries, Q.Z. conducted the experiments/analysis with the assistance of W.V. Q.Z. and W.V. wrote the manuscript.

## Methods

### Experimental Animals

Three rhesus monkeys (Macaca mulatta; 1 male; 4-7 kg) participated in this study. They were implanted with custom-built 8-channel phased-array receive coils (Janssens et al., 2012) embedded in an MRI-compatible head post prior to all procedures. Afterwards, they were trained using operant conditioning techniques to maintain fixation at a centrally presented fixation spot while their heads were constrained by the head posts in a natural ‘sphinx’ position inside a plastic box. Immediately before each fMRI scan session, the monkeys received an injection of monocrystalline iron oxide nanoparticle (MION, Feraheme, AMAG Pharmaceuticals; 8-11 mg/kg) through the femoral/saphenous vein to improve contrast-to-noise ratio (CNR) (Vanduffel et al., 2001). After all the fMRI sessions, an iron chelator was injected (Desferal, i.m, 1-1.5g per day, during 3-7 days) to remove the injected iron. Animal care and experimental procedures were performed in accordance with the National Institute of Health’s *Guide for the Care and Use of Laboratory Animal*, the European legislation (Directive 2010/63/EU) and were approved by the Ethical Committee of KU Leuven. Animal housing and handling were according to the recommendations of the Weatherall report, allowing extensive locomotor behavior, social interactions, and foraging. All animals were group-housed (cage size at least 16-32 m^3^) with cage enrichment (toys, foraging devices) at the primate facility of the KU Leuven Medical School.

### Apparatus and stimuli

The same stimuli as in Janssens et al. (2014) and Li et al. (2017) were used for the retinotopic mapping. A Barco LCD projector was used to project the stimuli at 1400 × 1050 resolution and 60 Hz refresh rate onto a translucent screen located 57 cm from the subjects’ eyes. Eccentricity and polar angle were mapped using phase-encoded annuli and wedges embedded with dynamic monkey faces (Zhu et al., 2012) and walking humans, respectively. Both the dilating and contracting annuli and the clockwise and counter-clockwise wedges were used to cancel phase errors caused by hemodynamic response delays. All stimuli traversed the area between 0.25° and 12.25° of visual angle in radius. Each run consisted of 4 stimulus cycles, and each cycle lasted 96 s. During a run, only one type of stimulus (e.g. wedge or annulus moving in one direction) was displayed. A central fixation point was presented constantly, and the subjects were required to maintain fixation on this point. In addition, hand positions were continuously monitored with an optic fiber and both hands had to remain in a rest position on two buttons in front of the monkeys in order to receive juice rewards-thus reward delivery was contingent on both accurate fixation behavior and fixed hand positions. An eye-tracking system based on infrared corneal reflection (ISCAN) was used to monitor subjects’ eye movements. Only runs in which the subjects maintained fixation within a small virtual fixation window (1.5° × 1.5° around the fixation point) for more than 90% of the time (and where the hands were in a rest position) were used for further analysis.

### Data collection

fMRI data were acquired at KU Leuven on a 3T Siemens TIM TRIO scanner with an AC88 gradient insert for M1 and M2 using the 8-channel implanted receive coils, and a custom-built local transmit coil. Submillimeter high-resolution T2* weighted EPI images (0.6 mm isotropic voxel size) were collected use a standard EPI sequence [repetition time (TR) = 3000 ms, echo time (TE) = 21 ms, acceleration factor = 2, flip angle (α) = 90°, acquisition matrix = 140 × 140 × 48]. Since only a limited number of slices could be scanned at this TR, data from the top and bottom parts of the brain of these two subjects were acquired from different sessions. fMRI data from M3 were acquired on a 3T Siemens PRISMA scanner with the implanted receive and local transmit coils, and a simultaneous multiple slice (SMS) EPI sequence (Nunes et al., 2006) was used to cover the whole brain [TR = 3000 ms, TE = 22 ms, accelerated multiband (MB) = 2, acceleration factor = 3, α = 90°, acquisition matrix = 140 × 140 × 74].

In addition to the EPI images, several T1 weighted (T1-w) 3D gradient echo (GRE) images (0.6 mm isotropic voxel size, TR = 6 ms, TE = 2.63 ms, α = 11°, matrix size = 160 × 160 × 96) were acquired in every fMRI session, to use as an intermediate image for the registration between functional and high-resolution anatomical images of the same subject.

High-resolution T1-w anatomical images of all subjects were acquired in separate sessions when the subjects were under ketamine-xylazine anesthesia. These images were used as the reference anatomy and to create a triangulated surface representation of the cortex for each subject. The anatomical images of M1 and M2 were acquired on the 3T Siemens TIM TRIO scanner using a single loop receive coil and the standard body transmit and gradient coils. 12 and 15 high-resolution T1-w images were acquired for M1 and M2 respectively, with a magnetization prepared rapid gradient echo (MPRAGE) sequence [0.4 mm isotropic voxel size, TR = 2500 ms, TE = 4.35 ms, inversion time (TI) = 850 ms, α = 9°, matrix size = 320 × 260 × 208]. The high-resolution anatomical images of M3 were acquired on the 3T Siemens PRISMA scanner using the same single loop receive, standard body transmit and gradient coils. 11 T1-w images were acquired with the same MPRAGE sequence (TR = 2700 ms, TE = 3.5 ms, α = 9°, TI = 882 ms, matrix size = 320 × 260 × 208).

To correct for EPI distortions caused by magnetic field inhomogeneity, high-resolution field maps (0.6 × 0.6 × 0.7 mm voxel size), consisting of two GRE images with different TEs, were acquired from the 3T Siemens PRISMA scanner for M3 during a separate session under anaesthesia (TR = 917 ms, TE_1_ = 6.48 ms, TE_2_ = 8.94 ms, α = 55°, matrix size 140 × 140 × 66). The same field maps were also acquired for subject M1 and M2, but in the same session as the experiments while the monkey was performing a fixation task.

### Surface reconstruction

To improve signal-to-noise ratio, several T1-w images from the same subject were averaged together to create the surface. Image segmentation and surface creation were performed using Freesurfer (Dale et al., 1999) following the similar procedure as described for humans. The submillimeter resolution was retained throughout all procedures.

### fMRI image reconstruction and pre-processing

To decrease artifacts caused by the movements of the monkeys, EPI images were reconstructed from the k-space using an “optimized generalized autocalibrating partially parallel acquisitions” (GRAPPA) reconstruction method (Hoge and Polimeni, 2015). The reconstructed EPI images were preprocessed using Freesurfer (http://surfer.nmr.mgh.harvard.edu) and Bioimage suite (http://bioimagesuite.yale.edu), through which skull stripping, slice-time correction, and motion correction was performed using one EPI volume as the template. The templates for M1 and M2 were one best EPI image for each monkey containing the least motion artifacts chosen from the best run where the monkey barely moved. For M3, the template was one EPI image acquired during the separate field map scanning session. Finally, a slice-by-slice undistortion correction was performed according to the template EPI using a B-spline grid-based nonlinear registration method to reduce frame-to-frame distortions (Li et al., 2017).

### GLM analysis

Polar angle and eccentricity maps were calculated following the procedure described by Sereno et al. (1995) from the preprocessed EPI images using Freesurfer. Nuisance regressors for the GLM analysis included three principal components derived from six motion-correction parameters and the linear and quadratic trend removals. The results were projected onto each monkey’s own surface to calculate field sign maps using the steps described below.

### EPI registration and surface projection

The same procedures as in our previous study (Li et al., 2017) were used to register each EPI template and the results in the same space to the reference anatomy of each subject. For M3, a distortion field was calculated based on the field map acquired in the same session as the EPI template and this was applied to the GLM results using FSL, thereby correcting for EPI distortions caused by magnetic field inhomogeneity. The resulting images were registered to M3’s reference anatomy through the intermediate 3D GRE image acquired in the same session using a rigid registration in Freesurfer. For M1 and M2, in which field maps were contaminated by susceptibility artifacts, a 1D nonlinear registration along the phase-encoding direction was performed using the JIP toolkit (http://www.nitrc.org/projects/jip) to calculate the distortions between the EPI templates and the GRE images. Furthermore, since fMRI and anatomical sessions in these two subjects were conducted using different gradient coils (AC88 gradient insert vs. standard gradient coil in the Tim Trio 3T scanner of Siemens), a gradient distortion correction described by Jovicich et al. (2006) was further performed to calculate the distortion field between each monkey’s GRE and their own reference anatomy images. The two distortion fields (EPI to GRE and GRE to anatomy) in each monkey were combined and applied to the GLM results before the results were registered to the reference anatomy through the intermediate 3D GRE image. The final results were then projected onto the surface, by only taking voxels located between 30% and 70% of the cortical depth between the grey-white matter boundary and pial surface. Values in these voxels were sampled in steps of 10% along the cortical thickness (depth sampling) and averaged to create the surface maps. This procedure not only minimized the effects of cerebrospinal fluid, white matter and pial vein signals on the maps but also reduced the cross-talk between neighboring superficial layers in the two banks within the same sulcus. Note that due to the use of a contrast image, large draining vein artifacts are absent (as opposed to standard BOLD imaging). No spatial smoothing was applied to the volume data. Heat kernel smoothing (bandwidth = 2, iteration = 10) (Chung et al., 2008) was applied to the surface data before iso-contour lines were calculated.

### Visual display of retinotopic maps

Unlike traditional maps from fMRI-based retinotopic studies, polar angle and eccentricity data are displayed as colored iso-contour lines on top of slightly transparent field sign maps (red and blue shadings) (e.g. Figure 2-4) (Sereno et al., 2015). This procedure allows visualization of visual field representation, local visual field sign and cortex structures (sulci and gyri, with dark- and light-gray shadings) on a single figure, in addition to fine-grained details of eccentricity and polar angle propagations. To improve visual segregation of UVF representations on these maps of the dorsal visual cortex where mainly the LVF is represented, we marked vertices with phase values > 0 (= UVF) with additional small black dots. Based on these contour and visual field sign maps, we manually drew summary diagrams using symbols for vertical and horizontal meridian, foveal and eccentricity representations (solid and dashed black thick lines, dashed white lines, yellow stars and purple thick lines). The vertical meridians were defined based on field sign reversals. The horizontal meridians were also based on field sign reversals in case the receptive field progression returned to the same quadrant, or between the upper and lower visual quadrant representations in case receptive fields crossed quadrants. It is important to note that in the former case, a 0° polar angle contour (corresponding to the HM) generally does not appear in fMRI data due to a coarse sampling and smoothness of the data: i.e. the HM is under-sampled relative to neighboring polar angle representations. In the latter case, the 0° contour does appear in fMRI data, however, it may not always perfectly align with the real HM presentation as illustrated by our simulations (see Supplemental Document and Figure S1). When the receptive field size is different on the two sides of the horizontal meridian, the 0° contour tends to shift towards the side with largest receptive field sizes (Figure S1A). The 0° contour also shifts towards the side with largest cortical magnification factors (Figure S1B). These situations occur at the anterior V2d border, near the gap in the non-mirror image visual field sign map (see white arrow in Figure 2A). Therefore, we drew the HM at that location slightly away from the 0° contour-line, which is fully supported by our simulations. This resulted in a continuous HM line parallel with the fundus of the lunate sulcus which followed the field map reversals along most of the border. It also resulted in a relatively fixed width of V2d throughout its entire extent, exactly as predicted from most anatomy-based definitions of V2d (Gattass et al., 1981; Sincich et al., 2003) (Figure 1). The foveal representations and eccentricity ridges were identified based on the (near)concentric iso-eccentricity lines and eccentricity reversals.

### Population receptive field size estimation

Population receptive field (pRF) size and position of each voxel in the grey matter ribbon of each subject was estimated based on the same submillimeter resolution retinotopic data using a model-fitting approach described by Dumoulin and Wandell (2008) and a MATLAB toolbox provided by Kay (2013). The pre-processed EPI data were first averaged across sessions and runs, independently for each stimulus type (dilating/contracting annuli, or clockwise/counter-clockwise wedges). Nuisance signals including 6 principal components derived from motion-correction parameters were removed from the EPI data before averaging. An equal number of runs were selected for each stimulus type, based on fixation performance. Runs with less than 90% fixation performance were discounted and the number of runs for each stimulus type was determined by the stimulus type with the least number of runs that survived that criterion in each subject. pRF sizes of dorsal and ventral areas in M1 and M2 were estimated separately from the data that were obtained in different sessions from the upper and lower part of the brain. The total number of runs selected for M1 and M2 for the two halves of the brain (top/bottom) were 30/11 and 17/11 per stimulus type. For M3, data from 13 runs were averaged for each stimulus type from the top and bottom part of the brain. A canonical gamma fit impulse response function (IRF) for the MION signal (Leite et al., 2002; Vanduffel et al., 2001) was used as the hemodynamic response function (HRF). The estimated receptive field size parameters (σ) were converted into degrees of visual angle and rendered onto the subjects’ surfaces. Values from 7 equally-spaced sectors (parallel to the cortical surface) covering the cortical tissue between the top 30% and bottom 70% of the cortical depth (between the pial surface and the grey-white matter boundary) were projected onto the surfaces separately, resulting in 7 surface maps. The estimated position parameters (x and y) were converted to complex values (magnitude set to 1) for polar angle and eccentricity representations and were projected onto the surfaces in the same way. The real and imaginary parts of these complex values were projected separately. To compare the retinotopic maps generated from the model-fitting approach with the GLM analysis, the real and imaginary components of the complex values were averaged across these 7 surface maps to generate polar angle, eccentricity, and field sign maps. The results of one representative subject are shown in Figure S4 and are very similar to the retinotopic maps generated using the GLM analysis.

### Region of interest (ROI) definition and pRF size extraction

The regions of interest (ROI) were drawn on the surfaces which were based on individual retinotopic maps. Vertices laying within ∼0.6 mm from the ROI borders were disregarded, to prevent cross-contamination of values sampled across different ROIs. The pRF sizes and eccentricities of all vertices from all 7 sectors parallel to the cortex (sampled from the top 30% to bottom 70% of the cortical thickness) and both hemispheres were pooled for each ROI in each subject. Vertices with poor pRF model fits (i.e. with explained variance < 35%) were excluded (Harvey and Dumoulin, 2011). Outlier vertices in each ROI were detected using a robust regression on the pRF size and eccentricity data. Vertices with a weight in the robust fit between pRF size and eccentricity exceeding 6 times the absolute deviation from the median weight were excluded. This removed less than 1.6% of data for each ROI across subjects. The pRF sizes in each ROI of each subject were then grouped into 48 equally spaced eccentricity bins spanning from 0.25 to 12.25 degrees of visual angle in log space. The median pRF size was calculated for each bin, and a robust regression was conducted on the median pRF size and eccentricity bins to further detect bins with extreme values using 3 times of absolute deviation as criteria. This removed 2.5 to 5.7% of data in each ROI.

### Quantification of pRF size differences across all sectors of interest in dorsal cortex

To statistically compare pRF size across all dorsal sectors with a quadrant representation, we fitted 3 linear mixed effect models with different complexities. All models included separate slope terms for the pRF size and eccentricity data from each area, using the maximum likelihood estimation in MATLAB and we compared the slopes across areas. The simplest model 1 was created to include a common intercept besides the slope terms as fixed effects and a single random effect corresponding to the intercept term to account for the subject random effects. The slightly more complex model 2 was created based on model 1, but included all the slope terms as well as the intercept in the random effects, to account for individual differences in slope. Model 3 was the most complex model, and was created by changing the common intercept term in model 2 in the fixed effects to separate intercept terms from different areas. The best-fitting model was selected using Schwartz’s Bayesian Information Criterion (BIC). The slope (fixed-effect) estimates across all ROIs were compared used a pairwise F-test and the P-values were corrected for multiple comparisons using a false discovery rate (FDR) correction including comparisons between all possible ROI pairs.

### Quantification of pRF size difference between R2 (DLP) and R1 (V3d)

To test whether foveal R2 (DLP) is more similar to its peripheral part as suggested by our retinotopic results, or more similar to R1 (V3d) as proposed by macaque models 1 and 3, we split R2 (DLP) into two parts using 2.5° eccentricity as cut-off and compared the pRF sizes between foveal and peripheral R2 (DLP) (comparison 1) or between R1 (V3d) and foveal R2 (DLP) (comparison 2). In both comparisons, V2d was included as a reference and split in the same way. In each comparison, a linear mixed effect analysis was performed in MATLAB and two linear mixed effect models were fitted to the data. In one model, a common intercept term but different slope terms were given to each of the two parts assumed to belong to the same area (in that comparison), e.g. foveal and peripheral R2 (DLP) in comparison 1, and the slope estimations between the two parts were compared. In another model, the slope term was fixed but the intercept terms were different for the two regions assumed to be the same area (in that comparison) and the intercept estimations were compared. In both models, random effects always included intercept and all slope terms. To test for an interaction effect, the difference between the two regions being tested [e.g. foveal and peripheral R2 (DLP) in comparison 1] were further compared with the difference between foveal and peripheral V2d using an F-test in MATLAB. The same analyses were also conducted using a much smaller cut-off eccentricity (e.g. 1°).

### Quantification of pRF size difference between R2 (DLP) and V3v

A similar linear mixed effect analysis was performed to compare the pRF sizes between V2v-V4v with their dorsal counterparts and R2- (DLP-), exactly as described above for the dorsal areas. A pair-wise comparison was conducted between all possible ROI pairs on the slope estimations using an F-test. The P-values were corrected for multiple comparisons using an FDR correction.

## Data and Software Availability

Data analyses were conducted using MATLAB (MathWorks), Freesurfer software suite (MGH), Bioimage suite (Yale University), analyzePRF (Kendrick Kay) and JIP toolkit (Joseph Mandeville). Software can be obtained upon request.

## Supplemental Information

Supplemental Document. Figures S1-S4.

### Supplemental Document: simulation details and results

#### Simulation I: Phase shift near the horizontal meridian (Figure S1)

Here, we simulated a few situations where phase shifts occur near the horizontal meridian (HM) between the upper and lower visual quadrant representations. In the first simulation, the receptive field sizes of the neurons representing the two quadrants were set to be different (Figure S1). In the second simulation, the cortical magnification factors (CMF) instead of the receptive field sizes between the two quadrants were different (Figure S1B).

In both cases, we simulated high resolution ‘neuron-like’ data by creating a 2D matrix with very high spatial resolution (504 × 504 matrix) to mimic a cortical sheet hosting the two neighboring areas. Each area occupied half of the matrix, as shown in Figure S1A, top left panel. The first area was located at the bottom half of the matrix and contained a lower visual field representation, in which receptive fields changed linearly from the lower vertical meridian (LVM, phase = -pi/2) to the HM (phase = 0). The second area was located in the top half of the matrix and contained an upper visual field representation, in which receptive fields changed linearly from the HM at the border shared with area 1 (indicated by the black dashed line in the middle of the matrix) to the upper vertical meridian (UVM, phase = pi/2).

In the first simulation, eccentricity phases increased linearly from the left to the right side of the matrix, thus from the most foveal (phase = 0) to the most peripheral representation (phase = 2*pi) in both areas. The slope of the linear change was set to be identical in the two areas, thus the CMF was the same. The same log transformation as used in the design of our stimuli for the retinotopy experiment was used to convert the eccentricity phase to the degree of visual angle, thus the CMF along the iso-polar angle direction approximated the organization of early visual areas (e.g. V1 and V2, etc.). The receptive field sizes of the two simulated areas were both set to be linearly related to eccentricity, but different slopes were assigned to the two areas, resulting in different receptive field size in the two areas. The formulas used for this simulation are:

Area 1: RFS = 0.24*E + 0.21

Area 2: RFS = 0.54*E + 0.21

where RFS is the receptive field size, and E is the eccentricity in degrees of visual angle. The top 3 panels in Figure S1A show the actual polar angle, eccentricity and receptive field size organizations that we designed for the ‘neuron-like’ data.

The simulated fMRI response evoked by the retinotopic stimuli used in our experiments from each of these neurons was generated using the analyzePRF toolbox (Kay et al., 2013). We used an impulse response function (IRF) for MION as the hemodynamic response function. To create the fMRI time course of each ‘neuron’ (i.e. cell in the matrix), the response was combined with 4 typical types of fMRI noise which were generated using an R package, neuRosim (Welvaert et al., 2011). The weight of each noise type in the nuisance signal was 0.1, 0.1, 0.3 and 0.5 for the white, temporal, physiological and spatial noise respectively. The baseline for the fMRI response was set to 1000, and the SNR was 40 when the 4 types of noise were added to the response. The gain parameter was set to 10 in the analyzePRF toolbox, and the peak value of the fMRI MION response before adding the noises was around −27 (0 was the baseline, note that we measure a negative instead of positive signal due to the contrast agent). Therefore, the percent signal change of the simulated fMRI response compared to the baseline (1000) was around 2.7%, which accurately reflects the real fMRI responses that we measured in the present experiments. Furthermore, we simulated the fixation performance by generating two temporal noise factors corresponding to eye movements in the horizontal (x) and vertical (y) directions, separately for each run. We restricted the fixation performance to be 95% within a fixation window similar in size, as we used in the experiment (1.5° × 1.5° of visual angle). The positions of the retinotopic stimuli were shifted around the center of the screen according to the simulated eye movements when the simulated fMRI responses were generated. The final fMRI time course data from the 504 × 504 ‘neurons’ were then downsampled to 28 × 28 voxels, with each voxel containing the response averaged from 18 × 18 ‘neurons’. The voxel data were then spatially smoothed with an FWHM of 3 and analyzed using the standard GLM procedure as in the real retinotopic experiment. Note that if the voxel size is 0.6 mm, as in our experiment, the width of each area will be 8.4 mm, which is very close to the width of V2 reported in the literature (Sincich et al., 2003). The results in 28 × 28 voxels space were up-sampled to 504 × 504, to enable comparisons (at the same resolution) with the initial (actual) polar angle and eccentricity representations of the models.

In the second simulation, we simulated the effect of CMF differences along the iso-eccentricity direction on the polar angle phase estimated at the horizontal meridian. To achieve that, the phase of the eccentricity across all the neurons in both areas was fixed to PI, which corresponded to 1.75° of visual angle, within the eccentricity range located at the gap immediately rostral to V2d (white arrow in Fig 2A). The receptive field size was fixed to 0.81°. Differences in CMF between the two areas were created by only adjusting the slope of the polar angle changes along the iso-eccentricity direction in area 2. This resulted in different width or different polar angle phases represented at the top border of area 2 if it was cut to the same width as area 1 as shown in the top panel of Figure S1B. In our case, the maximum phase at the top border of area 2 linearly changed from 0.18° to 90° from the left to the right side of the matrix. The fMRI time courses were simulated in the same way as described in the first simulation, and the data were down-sampled to the same number of voxels (28 × 28) to calculate the GLM results. The results were then upsampled to 504 × 504 resolution (Figure S1B, middle panel), from which the absolute 0 phase contour was calculated and the offset of this contour line in space was plotted in function of the maximum polar angle phase value represented at the top border of area 2, as shown in the bottom panel of Figure S1B.

#### Simulation II: pinched or narrow V3d (Figure S2)

In this simulation, we checked whether a pinched part of V3d (the first simulation, Figure S2A) or a very narrow part of V3d (the second simulation, Figure S2B) may exist at the gap neighboring V2d (white arrow of Figure 2A) without being resolved by the field sign analysis at 0.6 mm resolution (as in the actual experiments). In both simulations, a 512 × 512 neuron matrix was created to represent a cortical sheet hosting 3 areas. From the bottom to the top of the matrix, these corresponded to simulated areas V2d, V3d, and V3A. V2d was simulated to occupy a dimension of 256 rows × 512 columns, containing a lower visual field representation with the smallest receptive field size (RFS = 0.34*E + 0.21). V3A was simulated to occupy a dimension of 192 rows × 512 columns and covering an entire contralateral hemifield with the LVM shared with V3d, and with the largest receptive field size (RFS = 0.44*E + 0.21). In the first simulation, V3d was simulated to have a fixed narrow width (64 rows, which is ¼ of the width of V2d, as proposed for the pinched V3d, see macaque model 1 in Figure 1A). However, the phase of the polar angle represented at the border between V3d and V3A varied from −1° (-pi/180) to −90° (-pi/2). The goal was to identify the minimum polar angle phase represented at the top border of this pinched representation (analogous to the anterior border of V3d) to be detectable using the field sign analysis. In the second simulation, a LVM was represented at the anterior border of V3d, while the width of this area varied from 1 to 64 rows. This wedge-like V3d simulates a situation that enables us to identify the minimum width of V3d to be detected by the field sign analysis. In both situations, we tried to keep the eccentricities the same across all columns in all areas. However, since the gradient of the eccentricity needs to be calculated for the field sign analysis, we slightly varied the eccentricity in all areas from 1.44° (left) to 1.75° (right side of the matrix) of visual angle, which within the eccentricity range found at the gap neighboring V2d in our fMRI results. The receptive field sizes in V3d were simulated to be slightly larger than V2d, but smaller than V3A, as found in our results and previous studies. The formula we used was: RFS = 0.39*E + 0.21. The fMRI responses were generated in the same way as in Simulation I and the neuron data were downsampled to 28 × 28 voxels and spatially smoothed with an FWHM of 3 before the GLM and field sign analyses.

### Captions for Supplemental Figures

**Figure S1.**
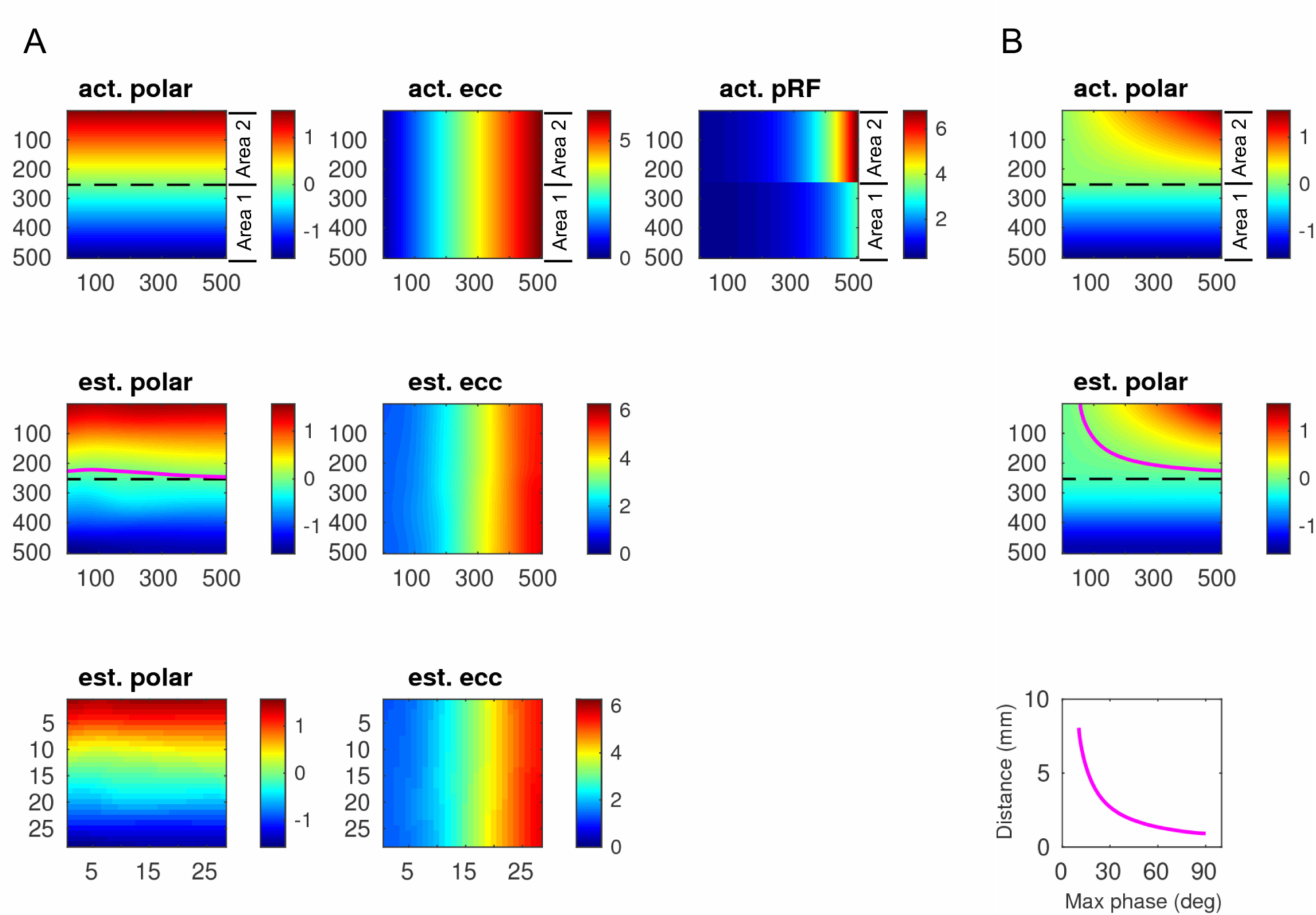
Results of Simulation I. (**A)** Different receptive field size between two neighboring areas shifts the polar angle phase at the horizontal meridian (HM) shared by them. Upper panels, the initially designed visual field representation; Lower panels, the estimated visual field representation from the GLM analysis from the down-sampled (simulated) fMRI data (28 × 28 voxels). Middle panels, GLM results in the lower panels upsampled to 504 × 504 resolution to compare with the initially simulated ‘ground truth’ data (shown in the upper panels). (**B)** Different cortical magnification factor along the iso-eccentricity direction shifts the phase at the HM. Upper panels: the initially designed visual field representation; middle panels, the estimated GLM results upsampled to 504 × 504 resolution; lower panels, offsets in space between the 0 phase contour line estimated from the GLM results and the real HM was plotted in function of the maximum polar angle phase represented at the top border of area 2. Black dashed line, real HM. White dashed line, estimated 0 phase contour line from the GLM results.

**Figure S2.**
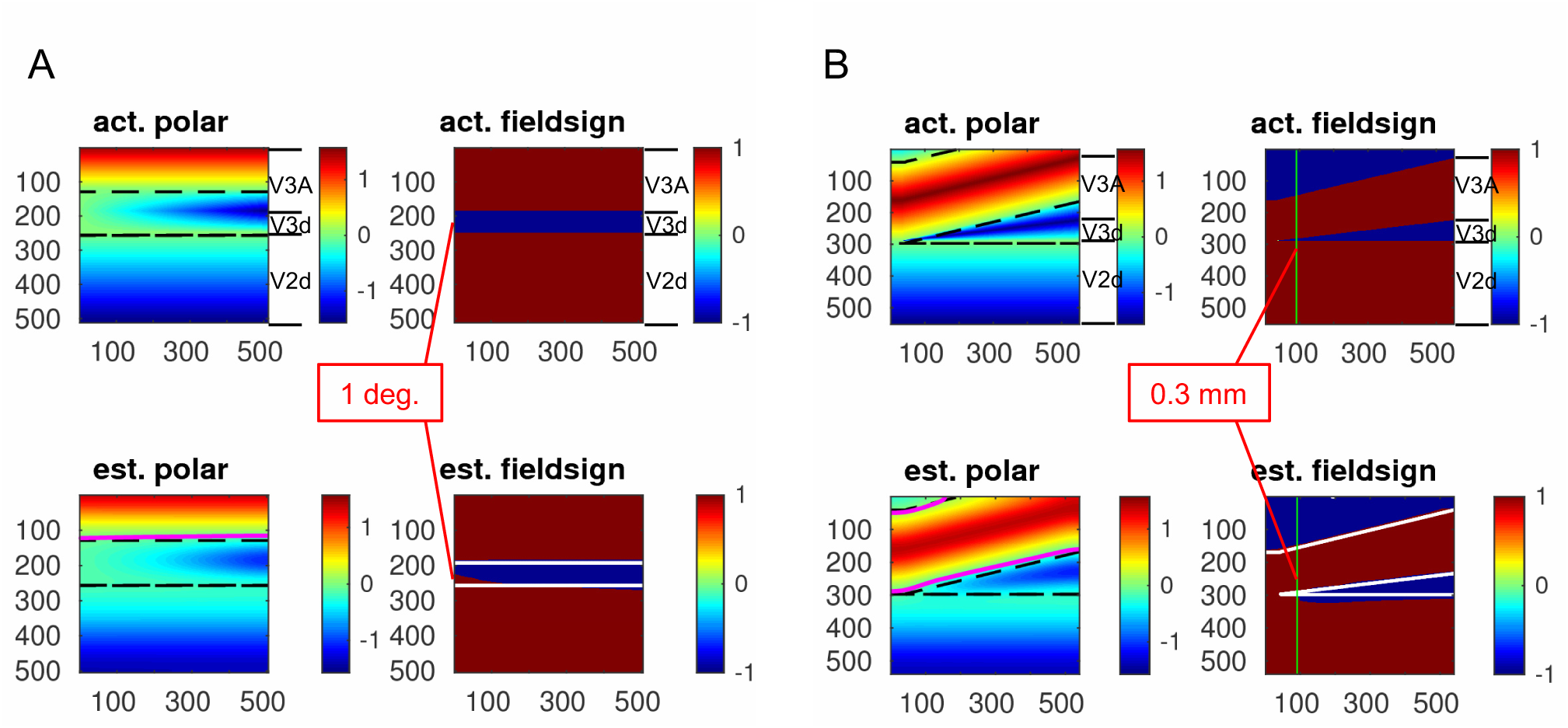
Results of Simulation II. (**A)** A pinched V3d with a polar angle as small as 1° represented at its anterior border could be detected by the field sign analysis at 0.6 mm resolution. (**B)** V3d needs to be smaller than 0.3 mm in width to remain undetected by our field sign analysis. The black dashed line (left panels in **A** and **B**) and white solid line (lower left panels in **A** and **B**) on the polar angle maps indicate the real and estimated 0 phase contour lines respectively. The white solid line on the estimated field sign maps on the right panels in **A** and **B** indicate the real areal boundaries as shown in the actual field sign maps.

**Figure S3.**
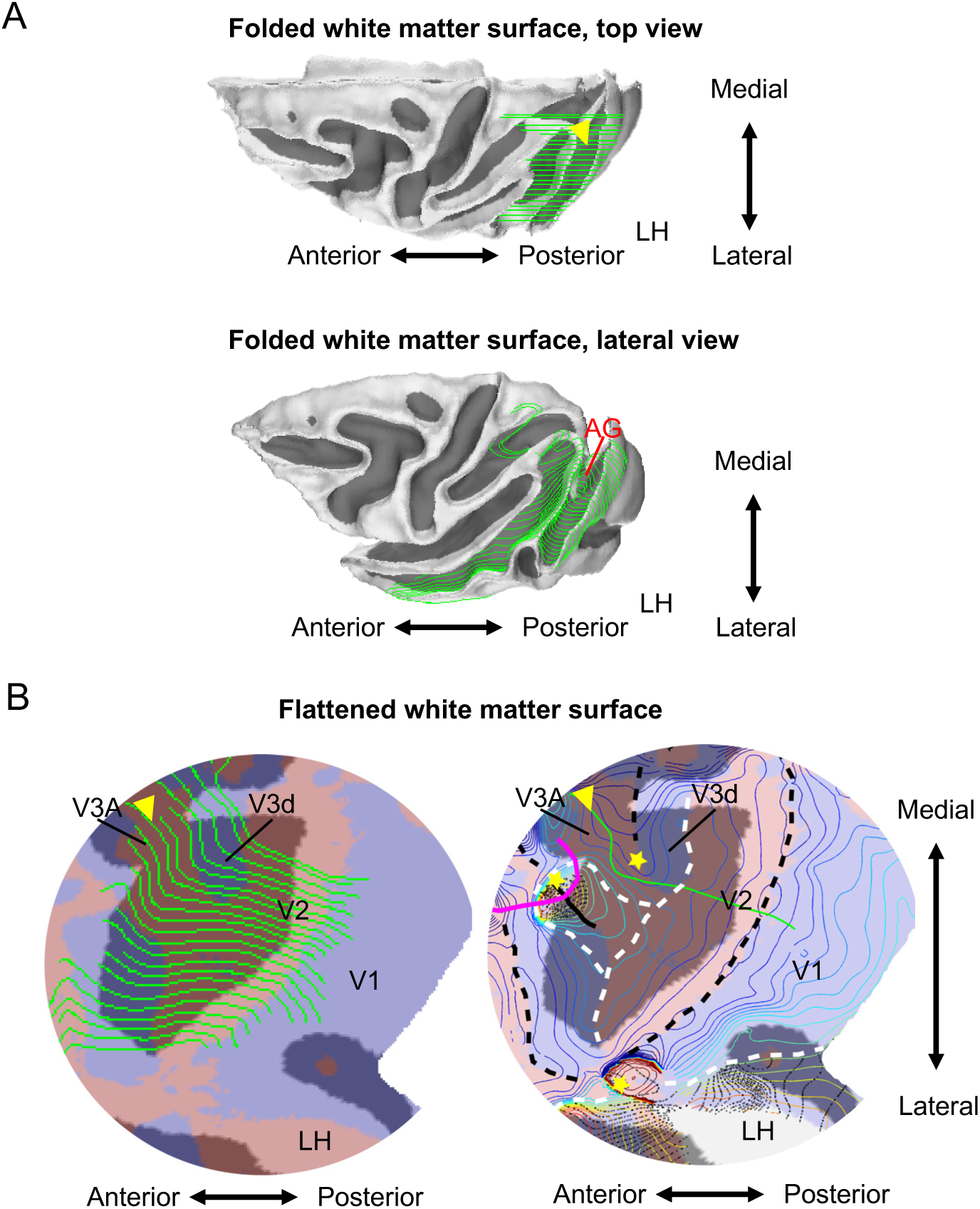
Recording sites of Nakamura and Colby on flatmap. Progression of recording sites along the annectant gyrus on sagittal sections as in Nakamura and Colby (2000; 2002) (**A**) can result in a distorted progression path running from lateral to medial on the flattened surface (**B**). The yellow arrowheads in **A** and **B** label the same path on the folded and flattened surface.

**Figure S4.**
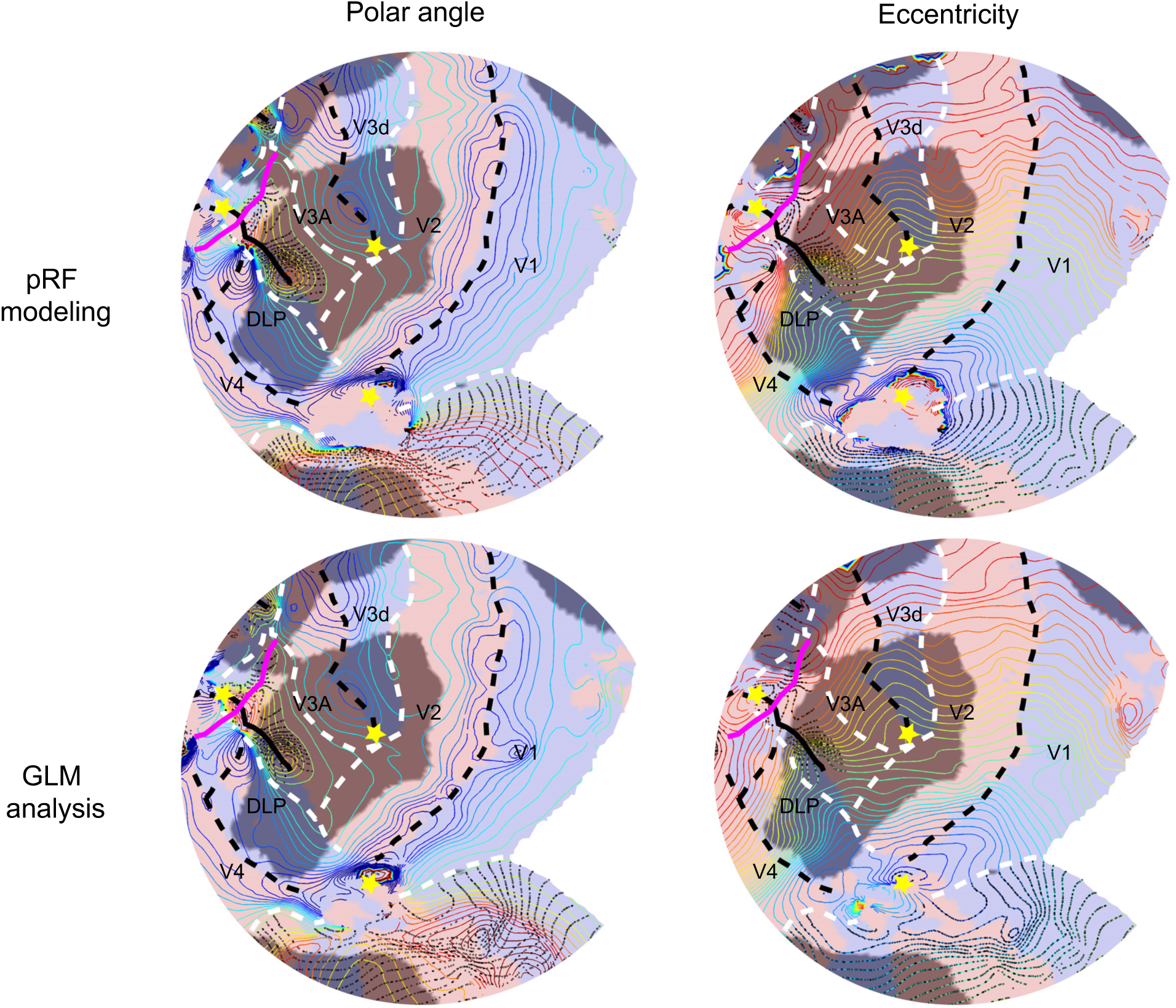
pRF- and GLM-based retinotopic maps. Retinotopic maps generated from the model-fitting (top panels) and GLM analysis (bottom panels) from M3. Note the excellent correspondence between the pRF- and GLM-based maps.

